# PKSmart: An Open-Source Computational Model to Predict *in vivo* Pharmacokinetics of Small Molecules

**DOI:** 10.1101/2024.02.02.578658

**Authors:** Srijit Seal, Maria-Anna Trapotsi, Vigneshwari Subramanian, Ola Spjuth, Nigel Greene, Andreas Bender

## Abstract

Drug exposure is a key contributor to the safety and efficacy of drugs. It can be defined using human pharmacokinetics (PK) parameters that affect the blood concentration profile of a drug, such as steady-state volume of distribution (VDss), total body clearance (CL), half-life (t½), fraction unbound in plasma (fu) and mean residence time (MRT). In this work, we used molecular structural fingerprints, physicochemical properties, and predicted animal PK data as features to model the human PK parameters VDss, CL, t½, fu and MRT for 1,283 unique compounds. First, we predicted animal PK parameters [VDss, CL, fu] for rats, dogs, and monkeys for 372 unique compounds using molecular structural fingerprints and physicochemical properties. Second, we used Morgan fingerprints, Mordred descriptors and predicted animal PK parameters in a hyperparameter-optimised Random Forest algorithm to predict human PK parameters. When validated using repeated nested cross-validation, human VDss was best predicted with an R^2^ of 0.55 and a Geometric Mean Fold Error (GMFE) of 2.09; CL with accuracies of R^2^=0.31 and GMFE=2.43, fu with R^2^=0.61 and GMFE=2.81, MRT with R^2^=0.28 and GMFE=2.49, and t½ with R^2^=0.31 and GMFE=2.46 for models combining Morgan fingerprints, Mordred descriptors and predicted animal PK parameters. We evaluated models with an external test set comprising 315 compounds for VDss (R^2^=0.33 and GMFE=2.58) and CL (R^2^=0.45 and GMFE=1.98). We compared our models with proprietary pharmacokinetic (PK) models from AstraZeneca and found that model predictions were similar with Pearson correlations ranging from 0.77-0.78 for human PK parameters of VDss and fu and 0.46-0.71 for animal (dog and rat) PK parameters of VDss, CL and fu. To the best of our knowledge, this is the first work that publicly releases PK models on par with industry-standard models. Early assessment and integration of predicted PK properties are crucial, such as in DMTA cycles, which is possible with models in this study based on the input of only chemical structures. We developed a webhosted application PKSmart (https://broad.io/PKSmart) which users can access using a web browser with all code also downloadable for local use.

Figure:
For TOC Only

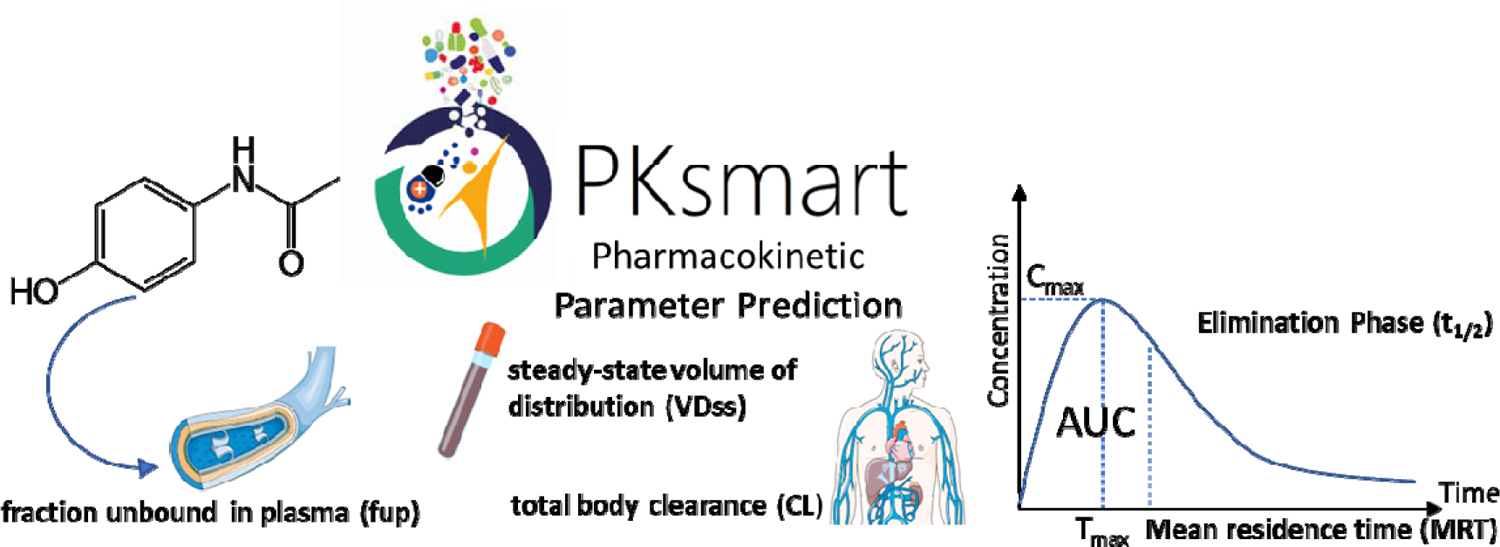

## INTRODUCTION

The mechanism of action of a compound, especially at an organism level, is not only dependent on the biological activity of the compound but also its exposure^1^, which can be defined by factors such as drug bioavailability, the volume of distribution, and clearance among other parameters.^2^ One way to estimate drug exposure *in vivo* is the extrapolation from *in vitro* data (also known as IVIVE: *in vitro* to *in vivo* extrapolation). However, predictions for *in vivo* intrinsic clearance tend to be underestimated for drugs with high observed *in vivo* intrinsic clearance.^3^ Other methods of estimating pharmacokinetics (PK) parameters include allometric scaling^4^ (or single-species scaling) and physiologically based pharmacokinetic modelling (PBPK)^5^. Previously, simple allometry and interspecies scaling^6^ have been used for the prediction of PK parameters, such as CL with an average fold error < 2.0 for small molecules.^7^ Allometric relations between rat and human CL, VDss and t½ have been reasonably accurate^8^, however sometimes have a higher error in estimation. For example, the prediction of volume of distribution in rats is prone to enterohepatic recirculation causing overestimation if allometrically scaled for the human volume of distribution.^9^

Commonly measured human PK parameters include the steady-state volume of distribution (VDss), clearance (CL), half-life (t½), fraction unbound in plasma (fu) and mean residence time (MRT). The VDss reveals the compound distribution between tissues and plasma, hence being dependent on both blood protein binding and tissue protein binding and is considered to be one of the least biased and one of the most reliable indicators of the extent of distribution.^10^ CL reveals the rate at which a drug is permanently removed from the plasma.^11^ The mechanisms of VDss are based on drug binding with tissue components while for CL, complex mechanisms such as metabolism and excretion via multiple pathways are involved.^11^ t½ represents the time taken for the drug concentration to reach half the initial concentration in plasma, while the MRT represents the average time spent by a drug molecule inside the *in vivo* system.^12,13^

More recently *in vivo* PK data has been modelled directly using 2D descriptors, ADME/PK properties as well as administered dose, as shown in Table 3. Studies have used chemical structural data to predict the volume of distribution^14,15^, the elimination half-life^16^, clearance^17^, human plasma protein binding^18,19,20^ and fraction unbound in plasma^21^. In particular, Schneckener *et al.* used predicted *in vitro*, physicochemical and ADME parameters and chemical structural data to classify oral bioavailability in rats with a balanced accuracy of 69.5%.^22^ Obrezanova *et al.* used various machine learning algorithms, including graph convolutional networks, that rely on features derived from chemical structures, ADME and physicochemical properties to predict rat *in vivo* PK parameters of clearance (R^2^ = 0.63) and bioavailability (R^2^ = 0.55).^23^ Conformal prediction has been used for human steady-state volume of distribution predictions, using a test set of 105 compounds, achieving a 2-fold error of 64%.^24^ Another recent study uses conformal prediction to achieve a mean prediction error between 1.4 to 4.8 for human PK parameters of fraction absorbed, oral bioavailability, half-life, unbound fraction in plasma, clearance, the volume of distribution and fraction excreted.^25^ Miljković et al established the first comprehensive protocol for the curation of human PK data and used chemical structure and administered dose for 1001 unique compounds to predict the volume of distribution in steady state and achieved an R^2^ = 0.47.^26^ These studies show that modelling *in vivo* PK parameters directly from chemical data is possible (as commonly used in predicting toxicity of molecules^27^), and this is also advantageous in the drug discovery cycle. Further, it has been shown that using predicted *in vivo* and *in vitro* data can improve early detection of drug-induced liver injury and that biological data was predictive of drug-induced cardiotoxicity.^28,29^ This is of further interest for the generative design of molecules^30^ where the early estimation of in vivo ADME parameters is of key interest for pharmaceutical research as it can be then used early on in design, for example, in design-make-test-analyse (DMTA) cycles to prioritise compound with commonly measured PK parameters.^31,32^

In this work, we present the first public *in vivo* PK model based on previously published datasets.^33,34^ The models used in our PKSmart tool integrated predicted animal PK parameters with structural and physicochemical parameters to model the human PK parameters of the steady-state volume of distribution VDss(L/kg), clearance CL (mL/min/kg), half-life t½ (h), fraction unbound in plasma fu (dimensionless) and mean residence time MRT (h). The models also provide an associated fold error estimate (and a range of predictions) which is dependent on the similarity of the compound to the chemical space of the training data. PKSmart (https://broad.io/PKSmart) is freely available for integration into any design environment with all code also downloadable for local use.

## METHODS

### Data Processing

#### (a) Human intravenous pharmacokinetic parameters

Human PK data was extracted from a dataset assembled by Lombardo et al which comprised intravenous (IV) pharmacokinetic data for 1,352 compounds.^33^ These parameters included steady-state volume of distribution VDss(L/kg), clearance CL (mL/min/kg), half-life t½ (h), fraction unbound in plasma fu (dimensionless) and mean residence time MRT (h). As a part of the data curation, compound SMILES were standardised which involved sanitization, normalisation, greatest fragment chooser, tautomer enumeration, and canonicalization as implemented in RDKit^35^. The standardisation process then protonates the molecule at pH 7.4 (using DimorphiteDL) by adding/removing protons to the molecule to mimic its state at the specified pH. For duplicate records with identical standardised SMILES, we used median values for each endpoint (otherwise mean if only two duplicate records were present). Finally, to remove large molecules, the exact molecular weight of the compounds was calculated, and compounds with an exact molecular weight greater than 1.5 standard deviations of the mean were filtered out. A decadic logarithm transformation was applied to all PK parameters except fu. This led to a dataset of 1283 unique compounds with 1249 VDss annotations, 1281 CL annotations, 1265 t½ annotations, 879 fu annotations and 1243 MRT annotations (henceforth referred to as the human dataset and provided as Supplementary Table S1; see Supplementary Figure S1 for distribution of data).

#### (b) Rat, Dog and Monkey pharmacokinetic parameters

Distribution at steady-state (VDss), clearance (CL) and fraction unbound in plasma (fu) for intravenous (IV) dosing were compiled from another dataset assembled by Lombardo et al which comprised 399 drugs.^34^ After standardisation of SMILES using the same pre-processing as above (including protonation at pH 7.4 and a molecular weight filter of 1.5 standard deviations of the mean of this dataset) and a decadic logarithm transformation applied on PK parameters except fup, this resulted in a 34.7% sparse dataset comprising 371 unique compounds (henceforth referred to as the animal dataset and provided as Supplementary Table S2; see Supplementary Figure S2 for distribution of data).

#### (c) External datasets

We first compiled compounds from the source of the animal PK dataset (Lombardo et al^34^) which also contained the human VDss for 17 drugs. In addition to this, we compiled data for 51 new drugs from the literature (FDA novel drug approvals for 2021 and 2022) with VDss, CL, fu and t½ annotations.^36^ For CL, we used a dataset from Yap et al^17^, who compiled total clearance in humans for 503 compounds from the literature. Out of these, we found 256 unique compounds with CL annotations that were not present in the training data used in this study (compared with standardised SMILES). Overall, we combined these datasets resulting in 315 unique compounds that were not present in the training data used in this study when compared with standardised SMILES (including protonation at pH 7.4). This dataset contained 315 unique compounds with 51 VDss annotations, 302 CL annotations, 34 fu annotations and 38 t½ annotations which was used as the external test set and is released as Supplementary Table S3. Additional annotations for MRT could not be identified in the literature and hence no external test set was available for this endpoint.

#### (d) DrugBank dataset

To evaluate the coverage of our datasets in the drug space, we obtained the chemical structural information (as InChI) of 2611 approved drugs (small molecules) from DrugBank (v5.1.9)^37^ as a reference point. Further, we annotated the drug molecules with the anatomical therapeutic chemical (ATC) classification codes from the KEGG DRUG Database resulting in a dataset of 1324 drugs with associated ATC classes. We standardised the SMILES (including protonation at pH 7.4) and removed outliers based on molecular weight distributions below 1.5 standard deviations of the mean molecular weight in this dataset. Finally, we obtained 2,304 unique molecular structures for the drug space.

### Structural and Physicochemical data

Morgan fingerprints of radius 2 and 2048 bits computed from standardised SMILES as implemented in RDKit were used as structural features. For physicochemical properties, we generated Mordred descriptors (at pH 7.4) as implemented in the Python package Mordred Error! Bookmark not defined..

### Feature selection

Feature selection was performed for the human, monkey, dog, and rat datasets separately. For the Mordred descriptors, first, we used the scikit-learn^38^ v1.1.1 variance threshold module to remove features having a low variance below a 0.05 threshold. Second, for all features, we calculated all pairwise correlations and removed one of each pair of features with pairwise correlations greater than 0.95. Finally, we removed feature columns if their minimum or maximum absolute value was greater than 15 (hence feature scaling was not required for the Random Forest algorithm). Hence, we obtained 162 Mordred descriptors for the human dataset, 218 Mordred descriptors for the monkey dataset, 203 Mordred descriptors for the dog dataset, and 205 Mordred descriptors for the rat dataset. Next, for Morgan fingerprints, we applied a variance threshold to prevent a model from fixating on less informative, minor features specific to a narrow chemical space of the dataset. By filtering out low-variance features, we ensure a model learns from more general and broadly applicable features across diverse chemical structures. To this effect, we used variance threshold to remove features having a low variance below a 0.05 threshold, resulting in fingerprints of length 152 bits for the human dataset, 207 bits for the monkey dataset, 203 bits for the dog dataset, and 205 bits for the rat dataset.

### Chemical Space Analysis

To assess the variability of chemical space of both the human dataset (1,283 unique compounds) and the animal dataset (371 unique compounds), we calculated the mean of 5-nearest neighbour Tanimoto similarity of each compound with the rest of the compounds in the respective datasets. Tanimoto similarity was calculated based on 2048-bit Morgan fingerprints.

To visualise the chemical space coverage of the models trained on the human dataset, we used a principal component analysis (as implemented in scikit-learn^38^ v1.1.1). For this, we removed binary variables and selected 84 of the Mordred descriptors that were continuous (out of the 162 Mordred descriptors selected for the human dataset). The same 84 descriptors were used to define the physicochemical space of the 1,283 unique compounds in the human dataset, the 371 unique compounds in the animal dataset and the 2,304 unique compounds in the approved drugs dataset (from DrugBank as a reference point).

### Linear regression analysis between observed animal and human PK parameters

We calculated the overlap of compounds between the human and animal datasets for which data was available. We compared compounds for which VDss, CL and fu data were available for all four organisms (human, monkey, dog, and rat). We used linear regression to determine the coefficient of determination between human and animal PK parameters.

### Model Training

#### (a) Training Models on Animal PK data

We trained individual Random Forest regressor models (as implemented in scikit-learn^38^ v1.1.1) for the three PK parameters from each of the monkey, dog, and rat datasets using the selected Morgan Fingerprint bits and selected Mordred descriptors, resulting in 9 models as shown in Figure 1 Step 1. Each endpoint was modelled using a 5-time repeated, 5-fold nested cross-validation. The data was split in the outer split into 5 folds, out of which, 4 were used to train a model and the other fold was used as the test set. For training the model we used a 4-fold cross-validation. The hyperparameters were optimised during cross-validation using a grid search (Supplementary Table S4 lists the parameter grid used to optimise the Random Forest models) and the results were evaluated on the remaining test fold. This was repeated for all 5 test folds comprising the entire data. The entire process was repeated 5 times to generate different splits of data resulting in 25 test folds and corresponding 25 models. We used the lowest geometric mean fold error (GMFE) from these 25 test folds to obtain the best-performing model. The parameters of the best-performing model were used to retrain the model on the entire training data which was used as the final model.

**Figure 1:**
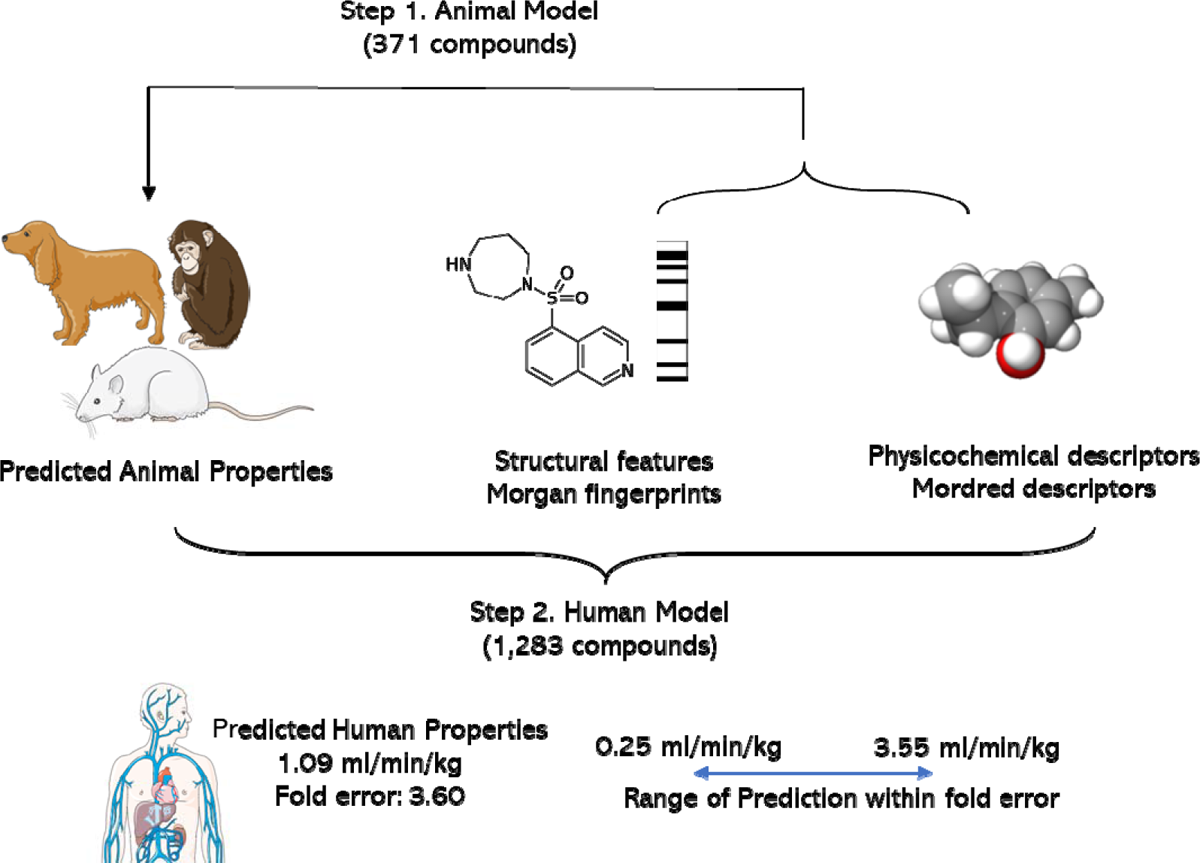
Workflow for models used in this study. First, models were trained on 371 compounds in the animal dataset to predicted animal PK parameters from structural fingerprints and Mordred descriptors. Second models were trained with different combinations of the real and predicted animal PK parameters with the structural fingerprints and Mordred descriptors for 1,283 compounds in the human dataset.

#### (b) Predicted Animal PK data for 1283 Human PK Dataset compounds

We used the 9 individual models trained to predict animal pharmacokinetic parameters of volume of distribution at steady-state (VDss), clearance (CL) and fraction unbound in plasma (fu) for all 1,283 compounds in the Human PK dataset as shown in Figure 1 Step 2. These predicted animal data were used as features for modelling the human PK parameters as described below.

#### (c) Training Models on Human PK data

For the five human PK parameters, the volume of distribution at steady-state (VDss), clearance (CL). half-life (t½), fraction unbound in plasma (fu) and mean residence time (MRT), we trained 9 types of Random Forest regressor models: (1) using Morgan fingerprints only, (2) and Mordred descriptors only, (3) using predicted animal data only, (4) using Morgan fingerprints and Mordred descriptors, (5) using Morgan fingerprints and predicted animal data, (6) using Mordred descriptors and predicted animal data, (7) using all three of Morgan fingerprints, Mordred descriptors and predicted animal data. In addition, we also built models (8) using a combination of predicted animal data and real animal data (where available) and (9) using Morgan fingerprints, Mordred descriptors and a combination of predicted animal data and real animal data where available. We compared these models to a baseline mean predictor model that always predicts the mean of the training data. This setup allowed us to compare different combinations of features to evaluate what combination is the best to predict human PK parameters.

For each human PK parameter endpoint and model combination, we followed the same procedure used previously to build models for animal PK parameters. We used a 5-time repeated, 5-fold nested cross-validation resulting in 25 test folds and corresponding 25 models from which we obtain the best-performing model with the lowest GMFE. The parameters of this best-performing model were used to train on the entire human dataset and this final model was used for predictions on the external test set.

#### (d) Calculating Fold errors

To calculate the fold error of a prediction for each endpoint, we looked at the trends for fold errors of predictions of all compounds as they appear in the 25 individual test sets. The structural similarity of each compound to their respective training data (in the particular iteration of the nested cross validation) was calculated as the mean Tanimoto similarity of 2048-bit Morgan fingerprints of the 5 nearest neighbours. For each value of structural similarity, we determined the mean fold error, and a kernel ridge regression (as implemented in scikit-learn^38^ v1.1.1) was fit on structural similarity to predict the mean fold error (we removed the compounds below 1.5 standard deviations of the mean similarity to training data). The kernel used was a combination of a radial basis function kernel and a white kernel to explain the noise of the signal that was optimised by a 10-fold grid search crossvalidation and a scoring function to maximise R^2^ (as implemented in Scipy v1.8.0^39^). Given the nature of structural similarity and mean fold error^40^, the RBF kernel may capture non-linear relationships while by including the white kernel, we account for that noise directly in the model simultaneously, aiming to get predictions that fit the data well but are not too sensitive to the noise. Finally, using this kernel, we estimate the fold error of prediction for a query compound in the final models: we fit the numerical value of structural similarity of this compound to the entire training data for the endpoint to the ridge regressor and assigned the predicted value as the fold error of the compound for the predicted endpoint. The Tanimoto similarity threshold of the test query compound was set at less than 0.25 which was outside 1.5 standard deviations of the mean of structural similarity to the training data of the final model and generally where the fold error tended to be greater than 5. If a test compound had a Tanimoto similarity less than 0.25 to the training data, an alert on the compound being outside the applicability domain of the model was raised along with the prediction and the estimated fold error. Besides the performance of the best models predicting human PK parameters on each of the individual 25 test folds of the nested cross-validation, we further evaluated our model on the predictions of human PK parameters for VDss, CL, and t½ for the external test.

### Comparison to in-house AstraZeneca Models

Proprietary pharmacokinetic (PK) models from AstraZeneca were procured, which included animal PK parameters for VDss, Cl, and fu for dogs and rats, and human PK parameters of VDss and fu. PK parameters from our study were extracted using the final PKSmart models which were developed in this study independently of the AstraZeneca models. We used Pearson correlation coefficients to assess the linear relationship between predictions from the AstraZeneca models and the predictions from the PKSmart models. The strength and direction of the correlations were interpreted to determine the comparability of the PKSmart models with the AstraZeneca in-house models.

### Model evaluation

We evaluated our models based on the 2-, 3and 5-fold error, median fold error (MFE), geometric mean fold error (GMFE) and bias which were used on the decadic antilogarithm transformation on the predicted values as defined by functions in released code. With regards to direct model performance measures, we used the root mean square error (RMSE) and coefficient of determination (R^2^) as implemented in the scikit-learn v1.1.1^38^, calculated by comparing the predicted values and the log transformed true values.

For a given predicted value *f_i_* compared to the true value *y_i_*, the fold error is given as:

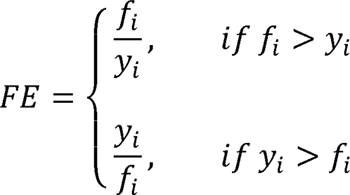

where i denotes an index running over n samples.

The 2-, 3and 5-fold error percentages were defined as the percentages of compounds for which the predicted value *f_i_* was within 2-, 3and 5-fold variabilities of the observed value *y_i_*.

The median fold error (MFE) and geometric mean fold error (GMFE) can be used to provide a measure of bias while considering equally all fold errors. The average logarithmic bias (ALB) is given as,

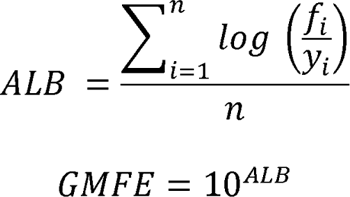

Another metric we used was the bias, which gives the median error between a predicted value *f_i_* and observed value *y_i_*.

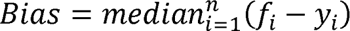

We also used metrics that consider individual prediction errors; we used the RMSE which measures the distribution of prediction errors.

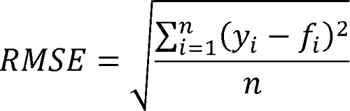

The coefficient of determination, R^2^ is defined as

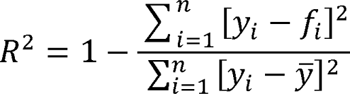

Where *y^—^* is the mean of the observed data

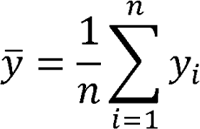

All code for these functions is released with the code on Github at https://github.com/srijitseal/PKSmart.

## RESULTS AND DISCUSSION

In this work, we built models to predict human PK parameters of volume of distribution at steadystate (VDss), clearance (CL), half-life (t½), fraction unbound in plasma (fu) and mean residence time (MRT) using a combination of Morgan fingerprints encoding structural information, Mordred descriptors encoding physicochemical properties and predicted animal PK parameters.

### Chemical Space Coverage

We first aimed to explore the chemical space in the human and animal datasets to evaluate the structural variance covered, where higher variance indicates a possibility to widen the applicability domain. As shown in Figure S3, we evaluated this using the distribution of the mean 5-nearest Neighbour Tanimoto similarity (using 2048-bit Morgan fingerprints) of each compound to the remaining compounds and found that 39.4% of compounds for the human dataset (and 49.3% of compounds for the animal dataset) lie below a 0.30 threshold of Tanimoto similarity to other compounds in respective datasets. This indicates that both datasets cover a wide range of structurally varying compounds, as shown by previous studies where 0.30 was deemed a plausible likelihood estimate of a threshold for similarity searching.^41,42^ Further, the compounds used in this study for the human and animal datasets represent a wide range of physicochemical properties as shown in Figure S4 for the descriptors of molecular weight (43.0 to 1,164.6 for the human dataset, 101.0 to 709.3 for the animal dataset), clogP (−16.6 to 11.4 for the human dataset, −11.29 to 5.78 for the animal dataset) and TPSA (0 to 569.1 for the human dataset, 4.44 to 338.41 for the animal dataset). As shown in Figure S5, the datasets cover a wide range of the relevant chemical space of DrugBank. Overall, this suggests that both the human and animal datasets are representative of a broad spectrum of the physicochemical space, enhancing their use in modelling PK parameters and broadening the applicability domain of the model.

While physicochemical descriptors capture certain properties of molecules, they do not encompass the entirety of a molecule’s biological activity or its interactions with biological systems. For this reason, we also compared the Anatomical Therapeutic Chemical (ATC) code distribution for these datasets. As shown in Figure S6, both the human dataset and the animal dataset covered a broad range of ATC code distribution at the top level (for 553 out of 1,283 compounds in the human dataset and for 235 out of 371 compounds in the animal dataset for which ATC annotations were available). This shows that the datasets encompass a vast array of approved drugs not only diverse in terms of chemical structures but also in their potential therapeutic applications.

### Distribution of PK parameters

We next analysed the distribution of the values for PK parameters in the human and animal datasets. Supplementary Figure S1 and S2 show the distribution of decadic logarithm-transformed data for each PK parameter and organism (human, dog, rat and monkey) combination, except fu for which the transformation was not applied. For human CL, out of 1,281 compounds, there were 1,180 compounds with a CL<= 25 mL/min/kg (low CL) and 101 compounds >25 mL/min/kg (high CL). Overall, most compounds (92.1%) in the human dataset exhibited low CL values which is often desirable for exposure but can lead to longer half-lives that are undesirable.^43^ On the whole, the datasets used in this study cover the diverse pharmacokinetic behaviour of compounds in different organisms.

### Animal PK parameters are predictive of human PK parameters

First, we analysed the animal PK data of VDss, CL and fu for their correlation to corresponding human PK parameters to evaluate translation from animal data to human data.^44^ As shown in Figure 2, we observe a linear relationship between human and monkey PK parameters (R^2^=0.74 with VDss for 91 compounds, R^2^=0.59 for CL for 95 compounds and, R^2^=0.53 for fu for 68 compounds) with similar trends observed for human vs dog and human vs rat PK parameters. While a correlation does not guarantee predictive accuracy, it does suggest that similarity in some physiological mechanisms. Previously, preclinical *in vivo* PK parameters from rat have shown to be advantageous for human PK prediction models.^45^ Given this potential similarity, we incorporated predicted animal PK parameters as an additional feature space for predicting human PK parameters.

**Figure 2:**
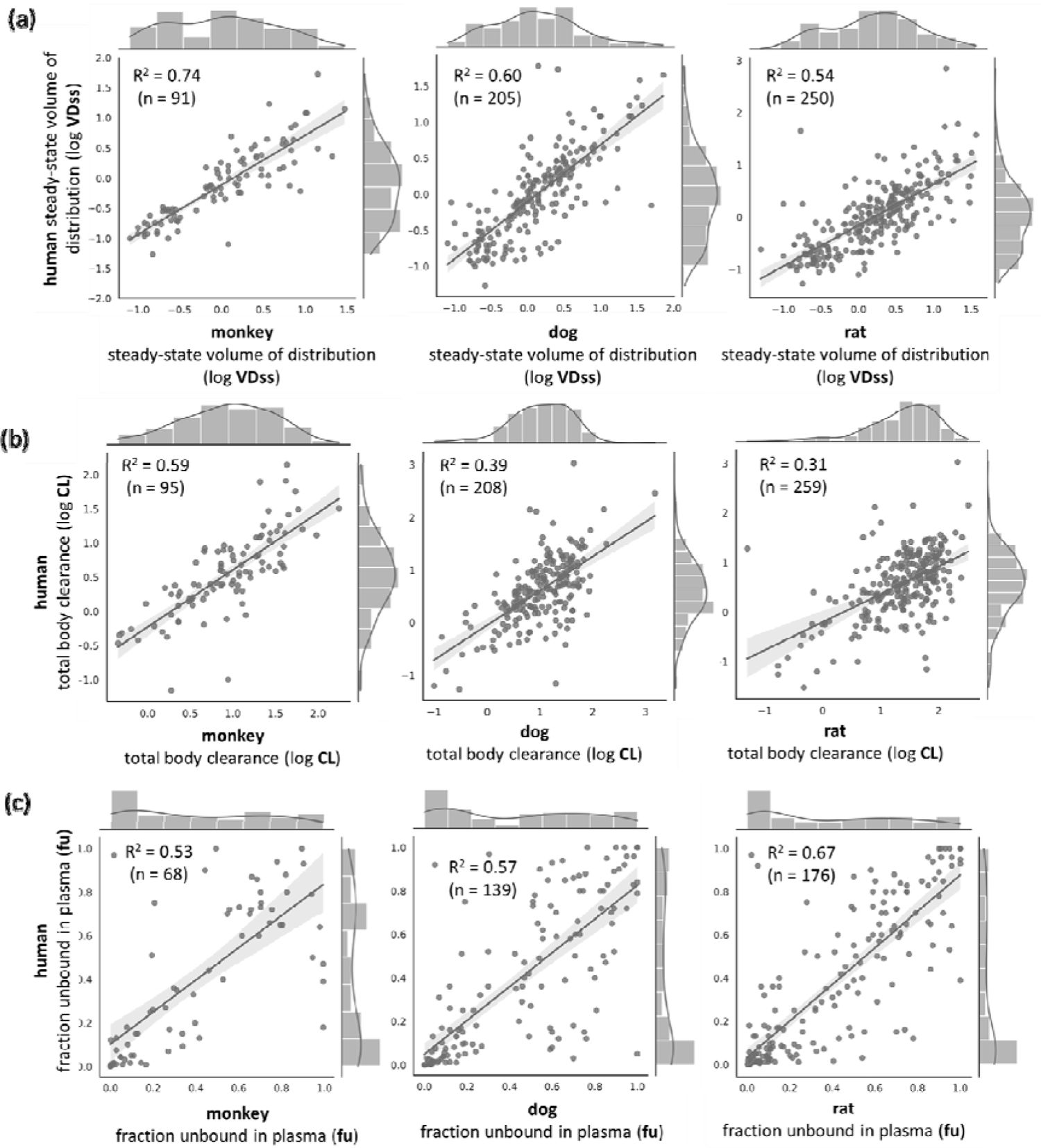
Linear regression fitting and coefficient of determination of the prediction of human PK parameters from animal PK parameters. Datasets were resampled such that the total number of unique compounds was the same for each endpoint of VDss, CL and fu.

### Structural and Physicochemical properties can reasonably predict Animal PK parameters

We trained nine individual models for VDss, CL and fu for each of the monkey, dog and rat datasets using Morgan fingerprints and Mordred descriptors in order to assess the possibility of modelling those endpoints using the chemical structure and physicochemical descriptors alone. Table 1 shows the median evaluation metrics of these animal PK models from the 25 test folds of the 5 times repeated 5-fold nested cross-validation. Overall, the best-predicted animal PK parameters were rat VDss (median R^2^ =0.46, RMSE= 0.43), monkey VDss (median R^2^ =0.46, RMSE= 0.44), dog CL (median R^2^ =0.29, RMSE= 0.40), monkey CL (median R^2^ =0.40, RMSE= 0.42), rat fu (median R^2^ =0.35, RMSE= 0.26), and dog CL (median R^2^ =0.43, RMSE= 0.25). We predicted the nine animal pharmacokinetic parameters for all 1,283 compounds in the human dataset and these were used as features for modelling the human PK parameters as described below.

**Table 1.** Median Evaluation metrics of animal PK models trained on Morgan fingerprints and Mordred descriptors from the 25 test folds of the 5 times repeated 5-fold nested cross-validation. Models predicted PK parameters for rats (324 compounds), dogs (264 compounds), and monkeys (128 compounds). GMFE: geometric mean fold error; RMSE: root mean square error.

| Organism | Endpoint | 2- fold<br>error<br>(%) | 3- fold<br>error<br>(%) | 5- fold<br>error<br>(%) | gmfe | bias | rmse | R<sup>2</sup> |
| --- | --- | --- | --- | --- | --- | --- | --- | --- |
| Rat | Clearance (CL) | 54.20 | 71.62 | 82.32 | 2.48 | -3.66 | 0.55 | 0.20 |
| Rat | Volume of<br>distribution (VDss) | 58.25 | 75.99 | 89.71 | 2.12 | 0.00 | 0.43 | 0.46 |
| Rat | Fraction unbound in<br>plasma (fu) | 57.30 | 70.70 | 79.07 | 2.72 | 0.03 | 0.26 | 0.35 |
| Dog | Clearance (CL) | 52.19 | 73.13 | 87.45 | 2.34 | -1.13 | 0.49 | 0.29 |
| Dog | Volume of<br>distribution (VDss) | 61.13 | 75.99 | 87.47 | 2.13 | 0.02 | 0.46 | 0.37 |
| Dog | Fraction unbound in<br>plasma (fu) | 62.68 | 74.79 | 82.69 | 2.50 | 0.02 | 0.25 | 0.43 |
| Monkey | Clearance (CL) | 53.21 | 80.08 | 92.58 | 2.12 | -0.56 | 0.42 | 0.40 |
| Monkey | Volume of<br>distribution (VDss) | 62.31 | 76.91 | 86.39 | 2.11 | 0.03 | 0.44 | 0.45 |
| Monkey | Fraction unbound in<br>plasma (fu) | 56.94 | 72.24 | 80.94 | 2.78 | 0.04 | 0.30 | 0.15 |

### Model predictions of Human PK parameters in a nested cross-validation

We trained 9 models to predict each of the human PK parameters VDss, CL, t½, fu and MRT as described in the Methods Section. It can be seen from Supplementary Figure S7 that 25 test folds were comparatively dissimilar compounds (<0.30 Tanimoto similarity) to the respective training data as shown by the distribution of the mean 5 nearest neighbour similarity of all test compounds over the 25 test folds in the nested cross-validation. The mean predictor was found to be the worst-performing model when considering median performance metrics over all 5 endpoints (as shown in Supplementary Table S5). We found that models using all three of Morgan fingerprints, Mordred descriptors and predicted animal PK properties (PKSmart models) achieved a higher median R^2^ (R^2^ = 0.55 for VDss, R^2^ = 0.31 for CL, R^2^ = 0.61 for fu, R^2^ = 0.28 for MRT and, R^2^ = 0.31 for t½) compared to models using only Morgan fingerprints (R^2^ = 0.46 for VDss, R^2^ = 0.24 for CL, R^2^ = 0.43 for fu, R^2^ = 0.25 for MRT and, R^2^ = 0.26 for t½) as shown in Figure 3. These models also improved 2fold errors marginally (from 55.2% to 58.0% for VDss, 48.4% to 51.4% for CL, 47.2% to 54.9% for fu, 48.6% to 50.4% for MRT and 47.8% to 51.0% for t½) compared to models using only Morgan fingerprints. As shown in Supplementary Table S5, similar trends of improvement were observed when models using all three of Morgan fingerprints, Mordred descriptors and predicted animal PK properties were compared to those using Mordred descriptors only or models using a combination of real and predicted animal PK parameters only. For CL and t½, models using Mordred descriptors and predicted animal data (CL: R^2^ = 0.31, t½: R^2^ = 0.31) and for CL, t½, and fu, models using Morgan fingerprints and predicted animal data (CL: R^2^ = 0.31, fu: R^2^ = 0.59, t½: R^2^ = 0.30) were sufficient for achieving the best performance with no significant improvement (as determined by a paired t-test as implemented in scikit-learn^38^ v1.1.1)) over models with all three of Morgan fingerprints, Mordred descriptors and predicted animal PK properties. Further, Figure 4 shows a significantly lower (using paired t-test) GMFE was achieved when using the models combining all three of Morgan fingerprints, Mordred descriptors and predicted animal PK properties (median GMFE = 2.09 for VDss, GMFE = 2.43 for CL, GMFE = 2.81 for fu, GMFE = 2.49 for MRT and, GMFE = 2.46 for t½) compared to other models using structural data using Morgan fingerprints only, Mordred descriptors only, a combination of Morgan fingerprints and Mordred descriptors. Therefore, the final model properties (PKSmart models) using all three feature spaces (Morgan fingerprints, Mordred descriptors and predicted animal PK properties) were the best-performing models to predict all five human PK parameters.

**Figure 3:**
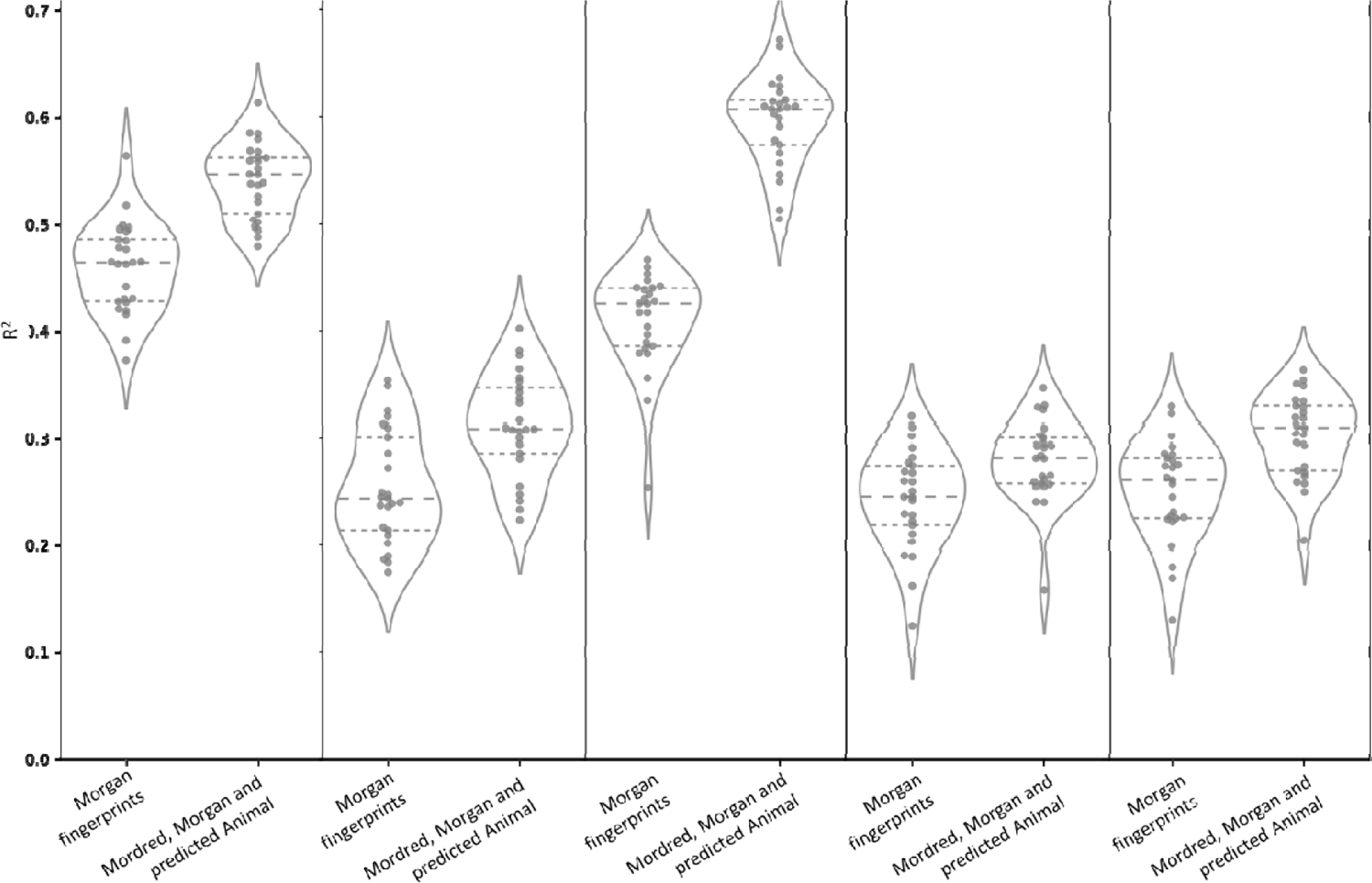
The distribution of coefficient of determination (R^2^) for models using Morgan Fingerprints versus models using a combination of Morgan Fingerprints, Mordred descriptors and predicted animal PK parameters over the 25 test folds in the nested cross-validation when predicting the five human PK parameters (a) VDss, (b) CL, (c) fu, (d) MRT and (e) t½

**Figure 4:**
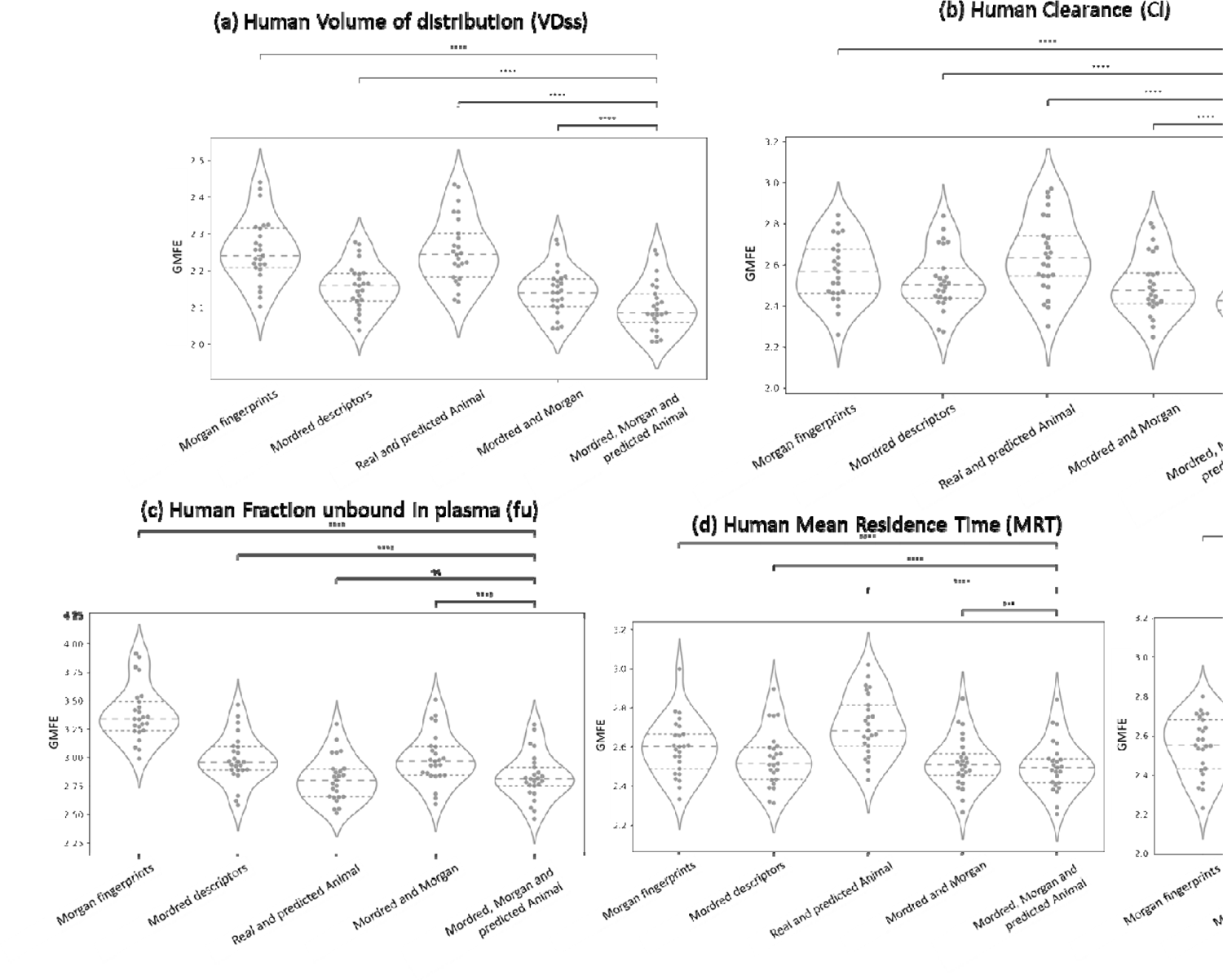
The distribution of geometric mean fold error (GMFE) over the 25 test folds in the neste acting the five human PK parameters (a) VDss, (b) CL, (c) fu, (d

### Evaluation based on Structural Similarity and Applicability Domain Considerations

We next looked at the mean fold errors for each compound that was structurally similar to the training data in the folds of the nested cross-validation for the model using a combination of Morgan fingerprints, Mordred descriptors and predicted animal PK properties. Figure 5 shows the kernel density estimate and Kernel ridge regression curves, which as expected show a decrease in fold error with increased structural similarity of a test compound to its respective training data. Further, we looked at the evaluation metrics using mean predictions for all compounds from all 25 folds of the repeated nested cross-validation as shown in Figure 6. It can be seen that there is a limited variance captured by the models which suggests that the models only capture some underlying signal in the training data. This is exemplified by Supplementary Figure S8 which shows the comparison of fold error of predictions to the respective observed human PK parameters, indicating fold errors tend to be large when the true value is further away from the majority of compounds (thus the observed value being out of the prediction interval of the Random Forest models). However, the models consistently demonstrated a lower RMSE than the baseline mean predictor as shown in Figure 6 (and details in Supplementary Table S5). Evaluation metrics were improved when applying applicability domain considerations (test compounds whose mean Tanimoto similarity to the training data is more than 0.30) for all five endpoints: VDss (R^2^=0.64, RMSE=0.39 for 731 compounds), CL (R^2^=0.45, RMSE=0.47 for 764 compounds, fu (R^2^=0.63, RMSE=0.21 for 501 compounds, MRT (R^2^=0.40, RMSE=0.47 for 730 compounds) and t½ (R^2^=0.42, RMSE=0.44 for 753 compounds). Additionally, considering the predictions based on the clearance classes of compounds suggests that 53.7% of the 650 compounds with low clearance and 59.5% of the 978 compounds with low to intermediate clearance**^Error!^ ^Bookmark^ ^not^ ^defined.^** (<12 ml/min/kg) were predicted to be within 2-fold error. However, only 10.4% of the 144 compounds with high clearance are within a 2-fold error range. The observed discrepancy in prediction accuracy across different clearance levels suggests inherent challenges in modelling compounds with high intrinsic clearance.^46^

**Figure 5:**
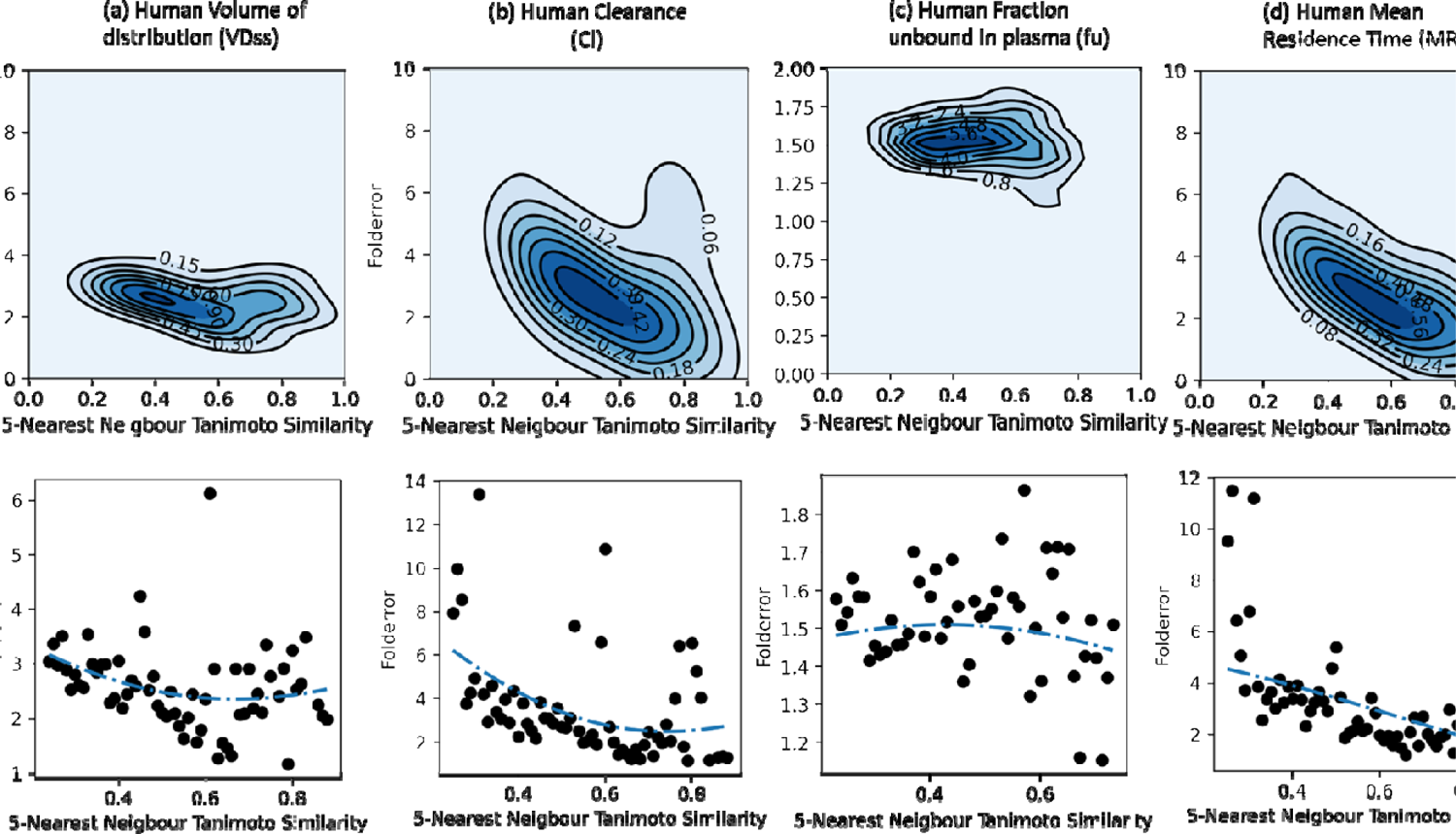
Kernel density estimate and Kernel ridge regression show a decrease in fold error with in the test compound to their respective training data during nested cross-validation for the five Dss, (b) CL, (c) fu, (d) MRT and (e) t½.

**Figure 6:**
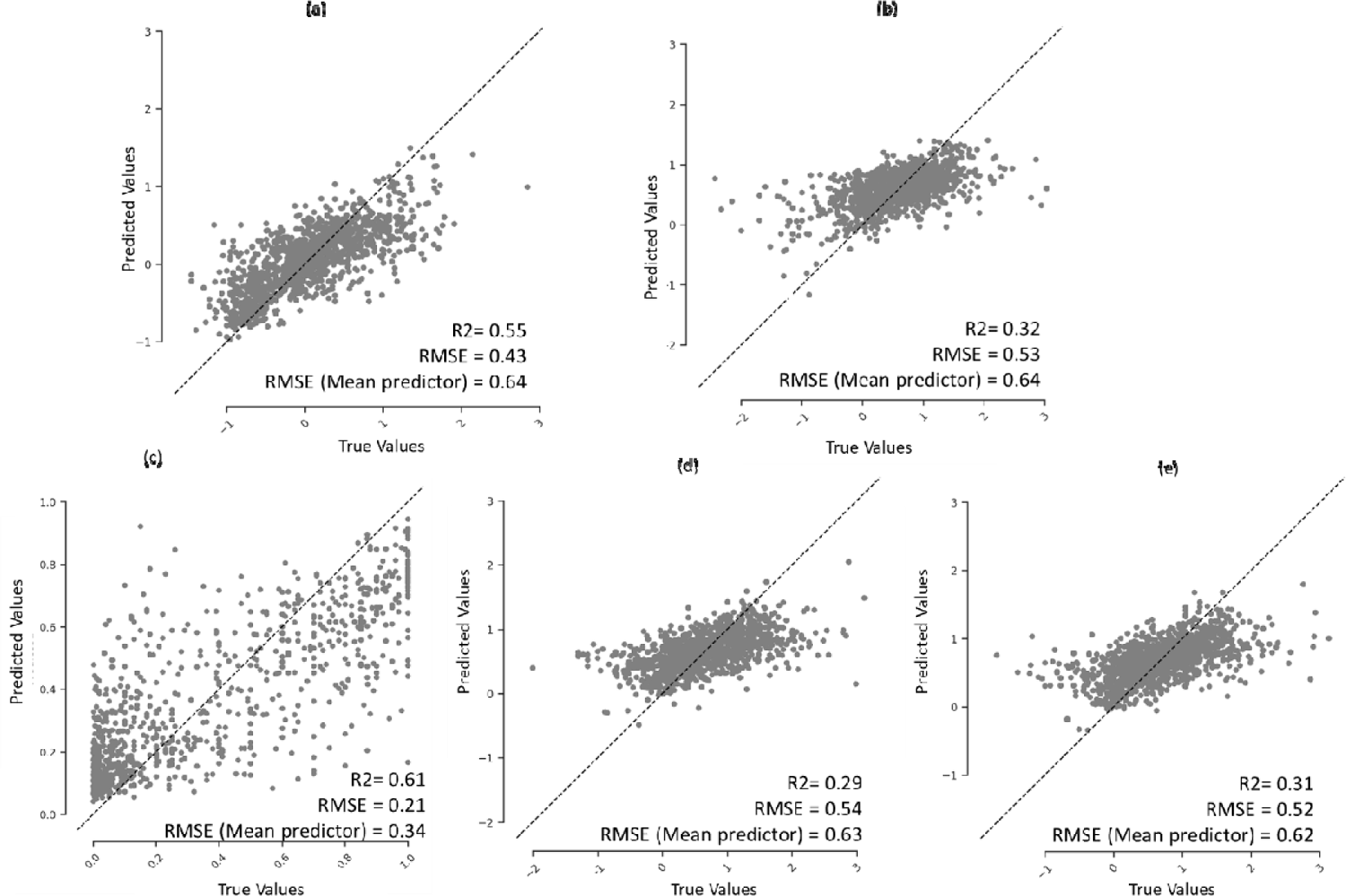
Regression plot of mean predicted PK parameters per compound over the 25 held-out test sets in the repeated nested cross-validation for the five human PK parameters (a) VDss, (b) CL, (c) fu, (d) MRT and (e) t½.

### Model Evaluation on the External Test Set

We next looked at the prediction compounds in the external test set that did not overlap with any of the unique compounds in the training data from the human dataset. Supplementary Figure S9 shows the pairwise Tanimoto similarity (and the contour graph) for 51 compounds for VDss, 302 compounds for CL, 34 compounds for fu, and 38 compounds for t½ in the external test dataset. The majority of pairs of compounds (over 99%) for all external datasets are structurally diverse with Tanimoto similarity <0.30. We compared models with all three of Morgan fingerprints, Mordred descriptors and predicted animal PK properties (as shown in Table 2) with evaluation metrics from both the repeated nested cross-validation and on the external test set (further details of individual predictions are shown in Supplementary Table S6 for all five PK parameters). The geometric mean fold error increased in the held-out test set compared to nested cross-validation for VDss (GMFE=2.58 in external test compared to GMFE=2.09 nested cross-validation), fu (GMFE=4.31 in external test compared to GMFE=2.81 nested cross-validation), and t½ (GMFE=3.31 in external test compared to GMFE=2.49 nested cross-validation) which can be attributed to the new chemical space. Nevertheless, the models remained reasonably accurate, with 43.6% of compounds within a 2-fold error and 62.2% within a 3-fold error for all four PK parameters. As shown in Table 2, predictions for CL had a lower GMFE =1.98 (R^2^=0.45) in the external test set compared to GMFE=2.43 (R^2^=0.31) in the repeated nested cross-validation.

**Table 2.** Evaluation metrics for 5 human PK properties on (a) nested cross-validation and (b) the external test set using models with all three of Morgan fingerprints, Mordred descriptors and predicted animal PK properties. GMFE: geometric mean fold error; RMSE: root mean square error.

| Endpoint | Method | 2- fold error (%) | 3- fold error (%) | 5- fold error (%) | gmfe | bias | rmse | R2 |
| --- | --- | --- | --- | --- | --- | --- | --- | --- |
| Volume of distribution (VDss) | Median from Repeated Nested-Cross-validation | 58.00 | 76.00 | 89.20 | 2.09 | 0.02 | 0.43 | 0.55 |
|  | Held-out-test (51 compounds) | 47.06 | 66.67 | 82.35 | 2.58 | -0.02 | 0.58 | 0.33 |
| Clearance (CL) | Median from Repeated Nested-Cross-validation | 51.36 | 71.09 | 84.77 | 2.43 | -0.22 | 0.53 | 0.31 |
|  | Held-out-test (302 compounds) | 66.89 | 79.80 | 87.09 | 1.98 | -0.03 | 0.45 | 0.45 |
| Fraction unbound in plasma (fu) | Median from Repeated Nested-Cross-validation | 54.86 | 66.48 | 77.27 | 2.81 | 0.04 | 0.22 | 0.61 |
|  | Held-out-test (34 compounds) | 23.53 | 47.06 | 55.88 | 4.31 | 0.07 | 0.23 | 0.23 |
| Half-life (thalf) | Median from Repeated Nested-Cross-validation | 50.40 | 70.28 | 83.47 | 2.49 | 0.08 | 0.54 | 0.28 |
|  | Held-out-test (39 compounds) | 36.84 | 55.26 | 73.68 | 3.31 | -3.66 | 0.69 | 0.05 |
| MRT | Median from Repeated Nested-Cross-validation | 50.99 | 71.54 | 85.38 | 2.46 | 0.18 | 0.52 | 0.31 |
|  | Held-out-test (0 compounds) | N/A | N/A | N/A | N/A | N/A | N/A | N/A |

### Comparison to in-house AstraZeneca Models

We compared models for human PK parameters using in-house AstraZeneca models (also built on the Lombardo dataset^33^) and the predictions from PKSmart models in this study. For both human VDss and fu, predictions from AstraZeneca models (R^2^ VDss: 0.30 and fu: 0.73) and PKSmart models (R^2^ VDss: 0.33 and fu: 0.21) are shown in Figure 8, which suggests that while PKSmart models are well predictive of VDss, they are not well predictive of the fu. As shown in Figure 7, when comparing the predictions for animal PK parameters for a set of compounds (where the experimental values are not known), we see a correlation in predictions for dog and rat PK parameters for both AstraZeneca and PKSmart models (Pearson Correlation R VDss: 0.67 for rat and 0.66 for dog, CL: 0.46 for rat and 0.49 for dog, and fu: 0.73 for rat and 0.62 for dog). This suggests that the PKSmart animal PK predictions correlate to the AstraZeneca’s internal models for VDss, fu and Cl animal PK parameters.

**Figure 7:**
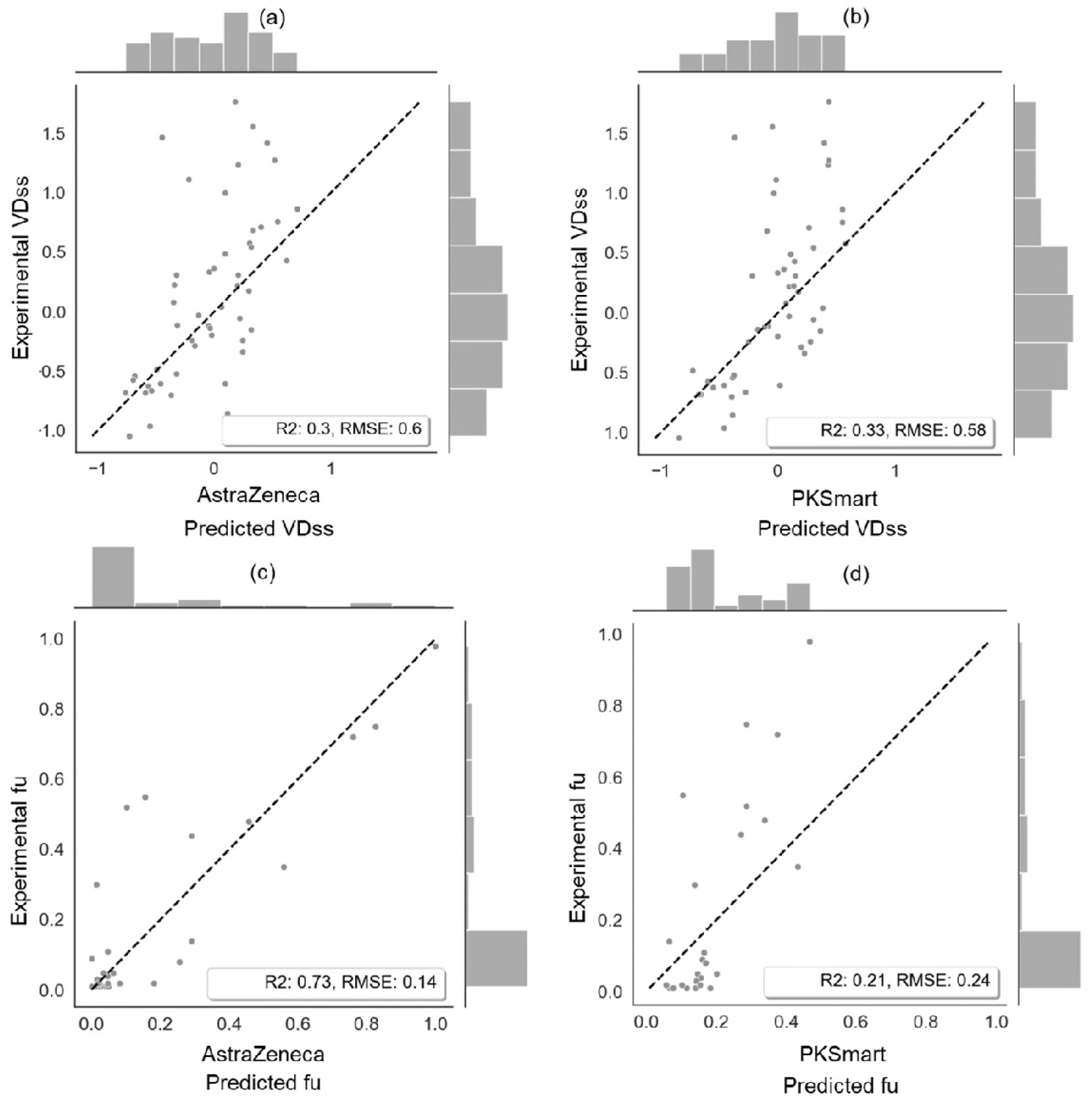
Performance of PKSmart model developed in this study and AstraZeneca model. Experimental values are plotted against the predictions for human PK parameter of (a) VDss using AstraZeneca model, (b) VDss using PKSmart model, (c) fu using AstraZeneca model, and (d) fu using PKSmart model.

**Figure 8:**
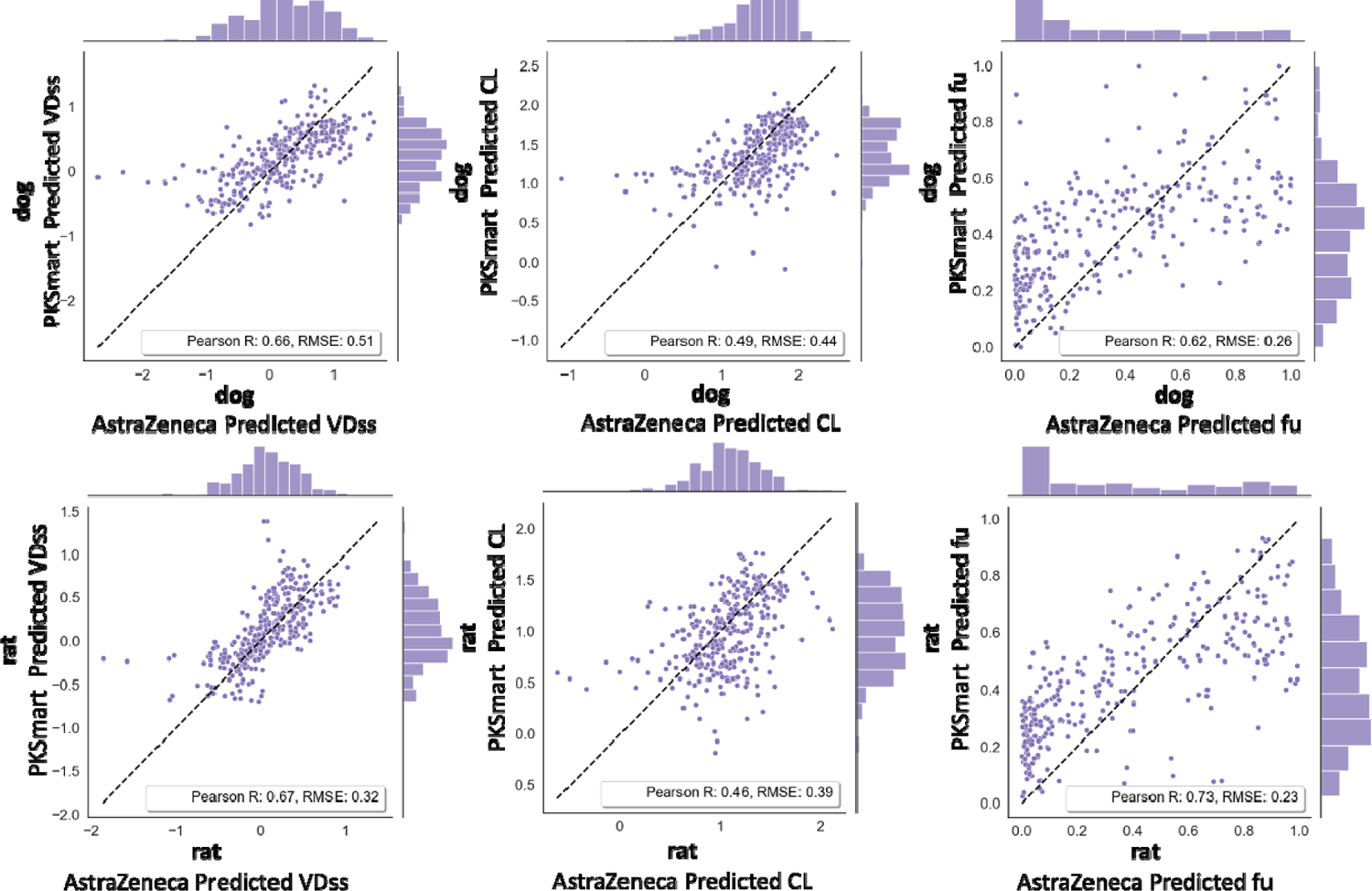
Correlation of PKSmart predictions versus AstraZeneca for animal PK parameters of rat (a) VDss, (b) CL and (c)fu, and dog (d) VDss, (e) CL and (f)fu for 315 compounds in the external test set.

### Comparison to Previous Literature

We next compared model performance to previously published literature. It needs to be kept in mind that models were established on very different datasets and validation methods and hence the metrics are not directly comparable. Table 3 shows the performance of some previously published PK models.^26,47,48,49^ Miljković et al predicted VDss using a curated dataset of 1001 unique compounds with an R^2^ of 0.47 (RMSE 0.50) for a held-out test set compared to this study where PKSmart models achieved an R^2^ of 0.55 (RMSE = 0.43) when using nested cross-validation on 1,249 compounds. Further the PKSmart models achieved an R^2^ of 0.33 (RMSE = 0.53) on an external test set of 51 compounds.^26^ Iwata et al used rat CL and chemical graph to model human CL to obtain GMFE of 2.68 (2-fold error of 48.5%) in cross-validation from 788 compounds compared to PKSmart models in this study which achieved a GMFE of 1.98 (2-fold error of 66.9%) when evaluated on an external test set of 302 compounds.^47^ While VDss was generally easier to model, the complex biology behind CL made modelling difficult for most published models. Hence the models developed in this study are at par with recently published literature.

**Table 3:** Evaluation metrics of previously published ML models compared to human PK models in this study trained on Morgan fingerprints, Mordred descriptors and predicted animal PK parameters. GMFE: geometric mean fold error; RMSE: root mean square error.

| PK Parameter | Source | Features | Compounds | Validation | R<sup>2</sup> | RMSE | GMFE | 2-fold error (%) |
| --- | --- | --- | --- | --- | --- | --- | --- | --- |
| CL (human) | Iwata et. al.<sup>47</sup> | Chemical Graph + rat CL | 748 | Cross Validation | - | - | 2.68 | 48.5 |
| CL (rat) | Kosugi et. al.<sup>49</sup> | Structural features and physicochemical descriptors | 1114 | Cross Validation | 0.56 |  |  | 71.9 |
| CL (human) | Wang et. al. | Molecular descriptors | 1268 | Cross Validation | 0.87 | 0.21 |  |  |
| CL (human) | Current Work | Morgan Fingerprints, Mordred Descriptors and Animal PK | 1281 | Median from Nested Cross Validation | 0.31 | 0.53 | 2.43 | 51.4 |
| CL (human) | Current Work | Morgan Fingerprints, Mordred Descriptors and Animal PK | 1281 | External Test set | 0.45 | 0.45 | 1.98 | 66.9 |
| VDss (human) | Miljković et. al.<sup>26</sup> | 2D, ADME/rat PK, Dose | 1001 | Held out test | 0.47 | 0.5 |  | 48.5 |
| VDss (human) | Current Work | Morgan Fingerprints, Mordred Descriptors and Animal PK | 1249 | Median from Nested Cross Validation | 0.55 | 0.43 | 2.09 | 59.0 |
| VDss (human) | Current Work | Morgan Fingerprints, Mordred Descriptors and Animal PK | 1249 | External Test set | 0.33 | 0.58 | 2.58 | 47.1 |

### Publicly available tool PKSmart

All code used to present results in this work is released publicly at https://github.com/srijitseal/PKSmart. All generated data from this code is further released at Zenodo (https://zenodo.org/doi/10.5281/zenodo.10611606).The final model that combined all three of Morgan fingerprints, Mordred descriptors and predicted animal PK properties was released as a python/streamlit-based web-hosted application PKSmart at https://pk-predictor.serve.scilifelab.se/ (also accessible via https://broad.io/PKSmart). Users can access the application using a web browser or locally with all code available via Zenodo at https://zenodo.org/doi/10.5281/zenodo.10611606.

## CONCLUSION

In this proof-of-concept, we used structural fingerprints, physicochemical descriptors, and predicted animal PK parameters to develop models for human PK parameters. We have developed the first publicly available tool using machine learning to predict these PK parameters. The web-hosted application developed in this study allows the user the predict the PK parameters from the input of chemical structure only and returns a range for each prediction with an estimated fold error of a compound based on the similarity to training data. This helps impart some understanding of the applicability domain of the models. Integrating animal PK features from across a range of species could therefore be used for fit-for-purpose and improved PK prediction in drug discovery. Such models can then be integrated into DMTA cycles to facilitate compounds with desirable PK parameters in the early stages of drug discovery. In the future, greater availability of public data could significantly improve predictive models. The web-hosted application PKSmart can be accessed at https://broad.io/PKSmart via web browser and with all code downloadable for local use at https://zenodo.org/doi/10.5281/zenodo.10611606.

## ASSOCIATED CONTENT SUPPORTING INFORMATION

We released the Python code for our models which are publicly available at https://github.com/srijitseal/PKSmart and code ready for local implementation is available via Zenodo at https://zenodo.org/doi/10.5281/zenodo.10611606. PKSmart is freely available at https://pk-predictor.serve.scilifelab.se (also accessible via https://broad.io/PKSmart).

## AUTHOR INFORMATION

All authors have approved the final version of the manuscript.

## ACKNOWLEDGMENT

S.S. acknowledges funding from the Cambridge Commonwealth, European and International Trust, Boak Student Support Fund (Clare Hall), Jawaharlal Nehru Memorial Fund, Allen, Meek and Read Fund, and Trinity Henry Barlow (Trinity College). S.S. acknowledges support with funding from the Cambridge Centre for Data-Driven Discovery and Accelerate Programme for Scientific Discovery under the project title “Theoretical, Scientific, and Philosophical Perspectives on Biological Understanding in the Age of Artificial Intelligence”, made possible by a donation from Schmidt Futures. All authors thank Erik Gawehn for discussing results and proof-reading the manuscript.

