## Supplementary Figures for "PKSmart: An Open-Source Computational Model to Predict *in vivo* Pharmacokinetics of Small Molecules"

Machine Learning, Toxicity, Bioactivity, Applicability Domain, Pharmacokinetic Parameters

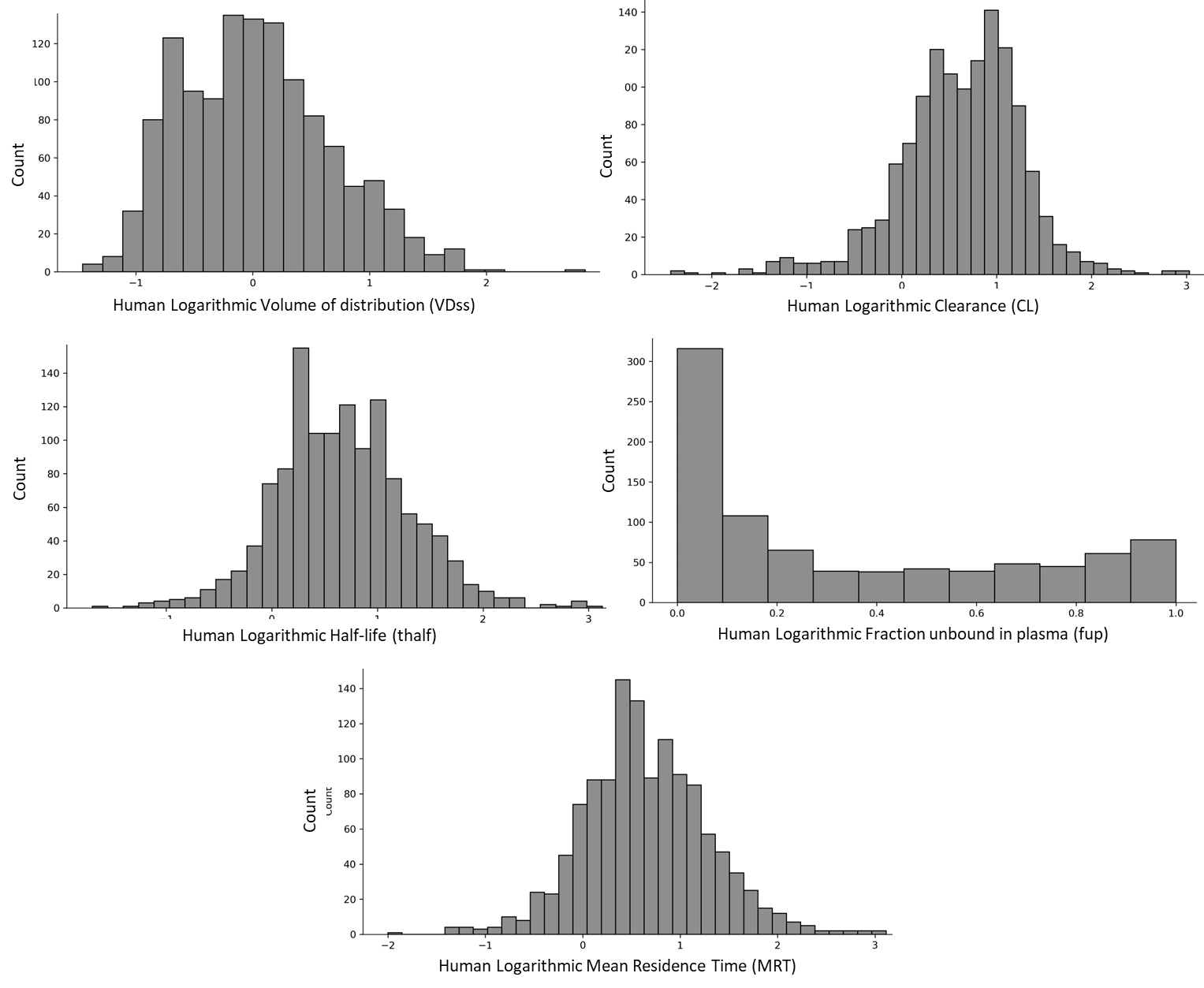

Figure S1: Distribution of 1,283 unique compounds with 1249 VDss annotations, 1281 CL annotations, 1265 t½ annotations, 879 fu annotations and 1243 MRT annotations in the human dataset.

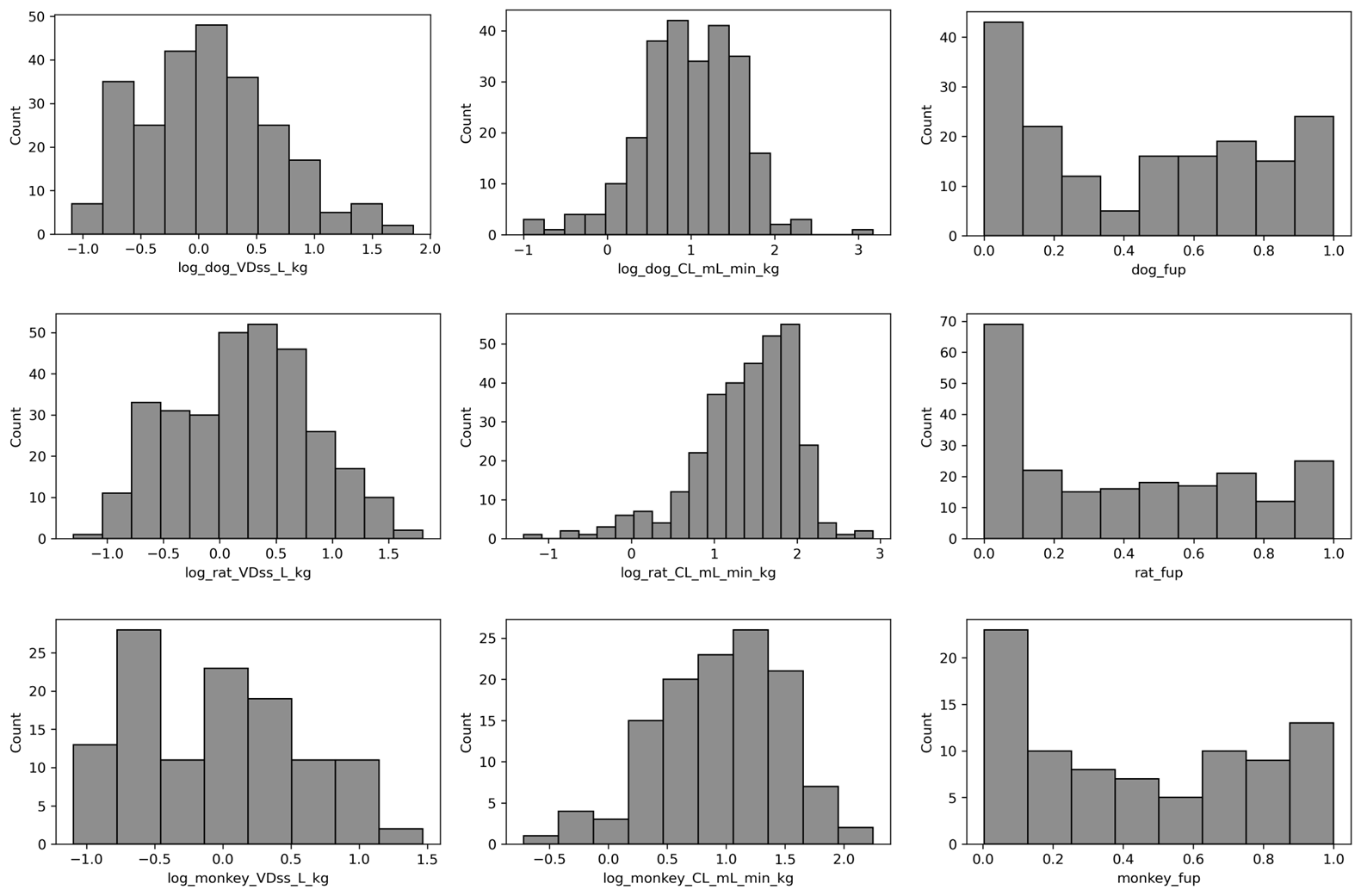

Figure S2: Distribution of 371 unique compounds with VDss , CL and fu annotations in the animal dataset for dog, rat and monkey.

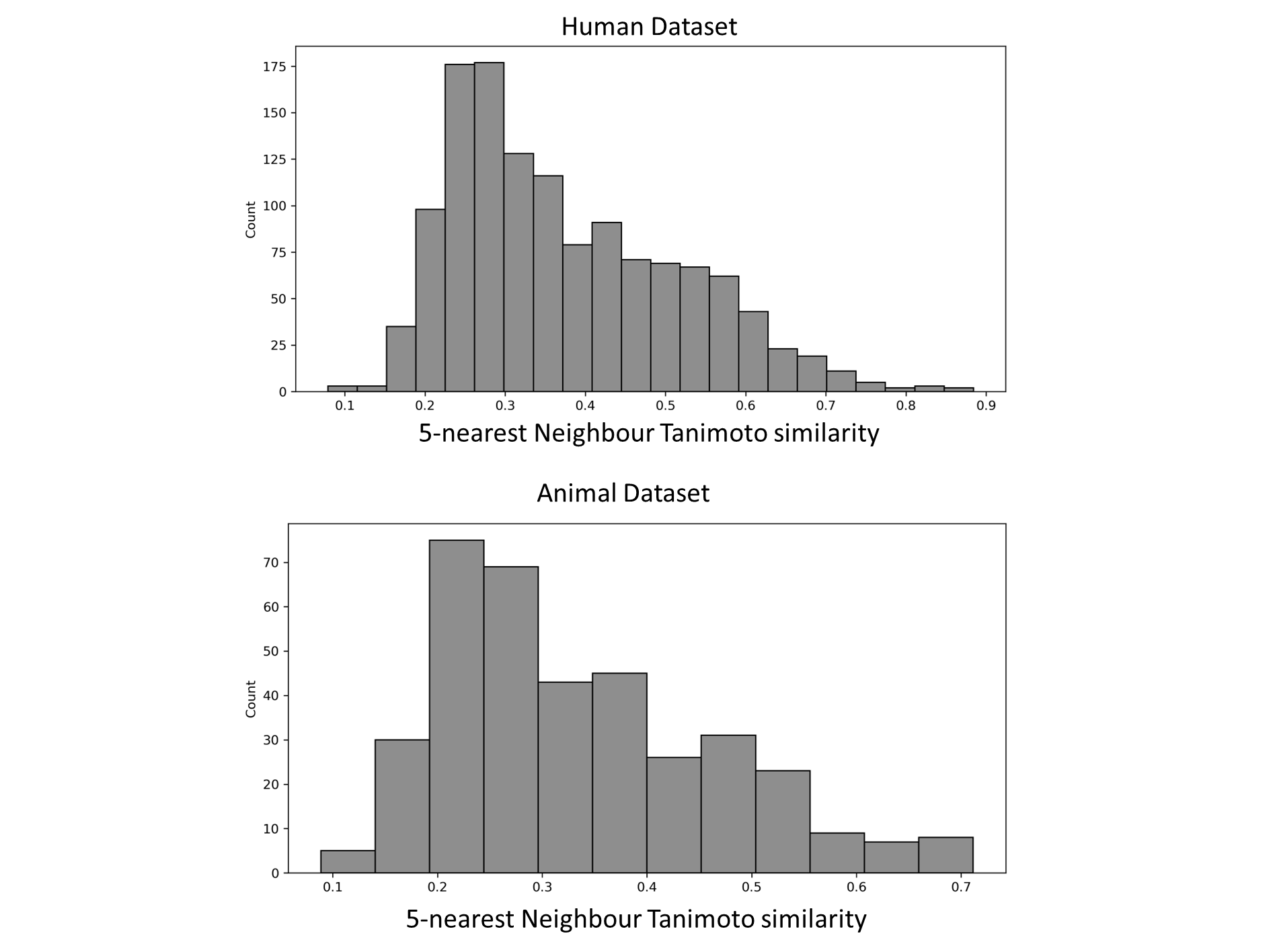

Figure S3: Distribution of the mean 5-nearest Neighbour Tanimoto similarity (using 2048-bit Morgan fingerprints) for each compound in the human and animal dataset compared to the other compounds in the respective datasets.

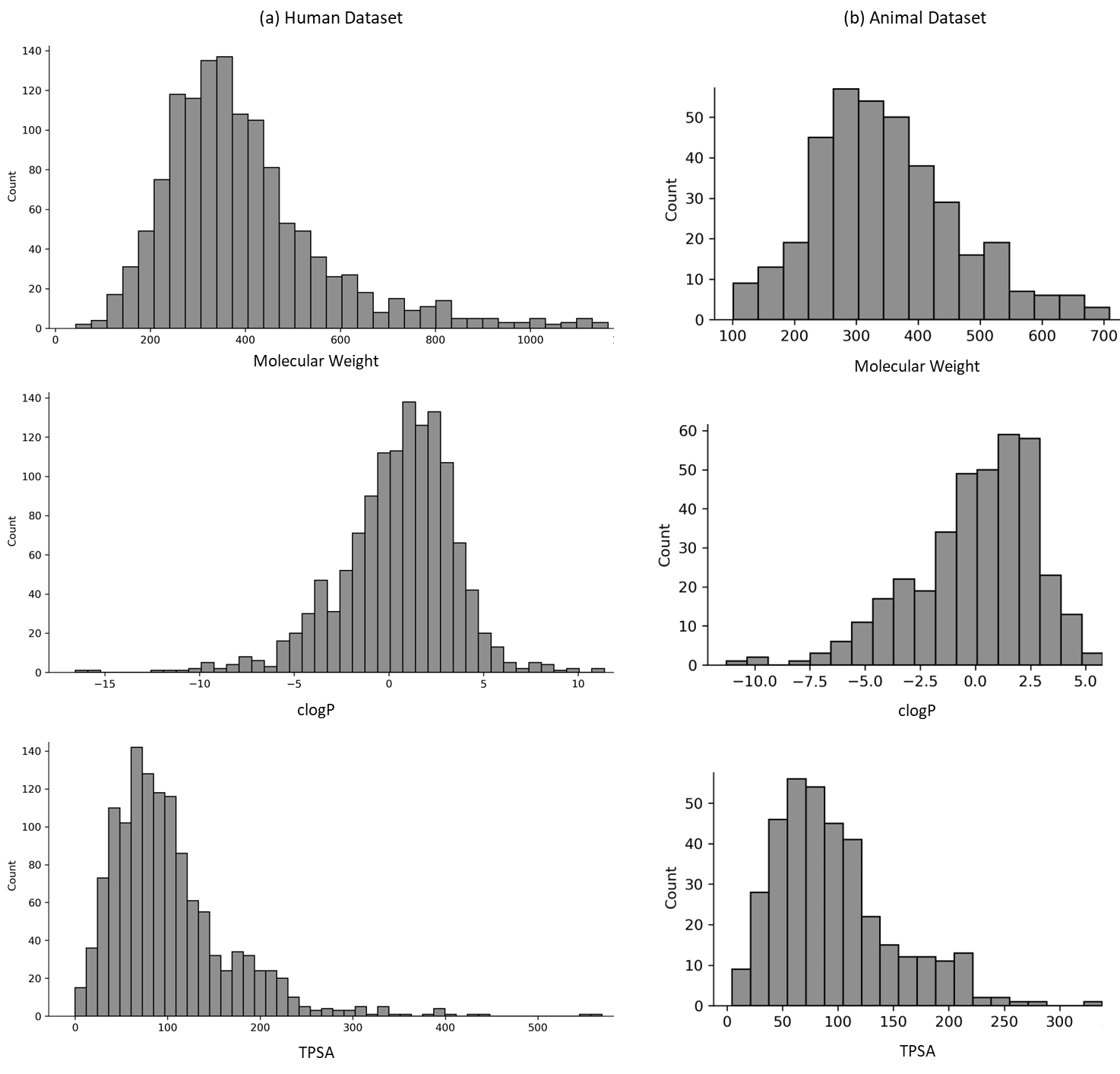

Figure S4: Distribution of physicochemical properties of molecular weight, clogP and TPSA for the human and animal dataset.

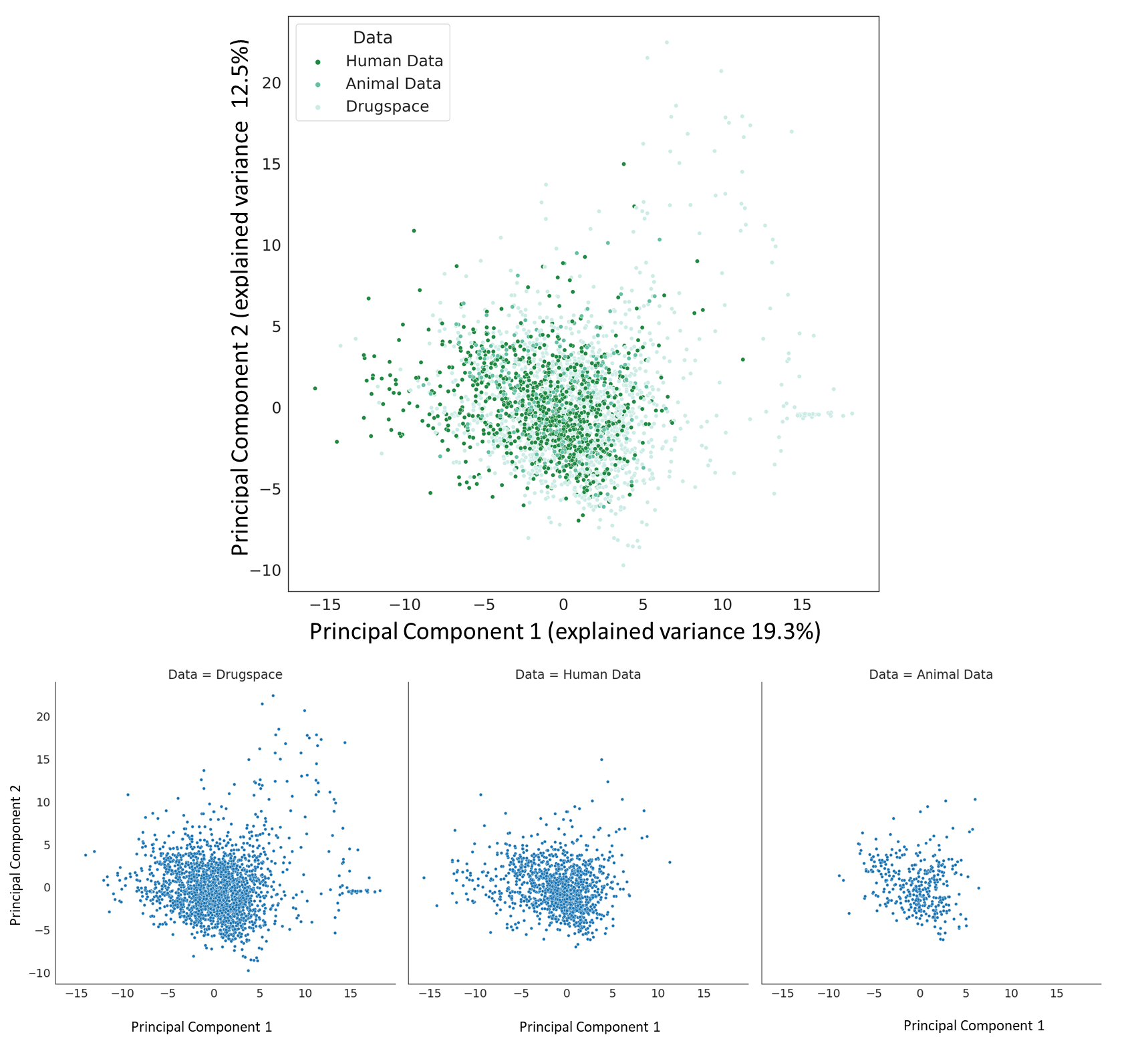

Figure S5: Principal component analysis shows the physicochemical space of the human and animal dataset overlayed on the physicochemical space of approved drugs.

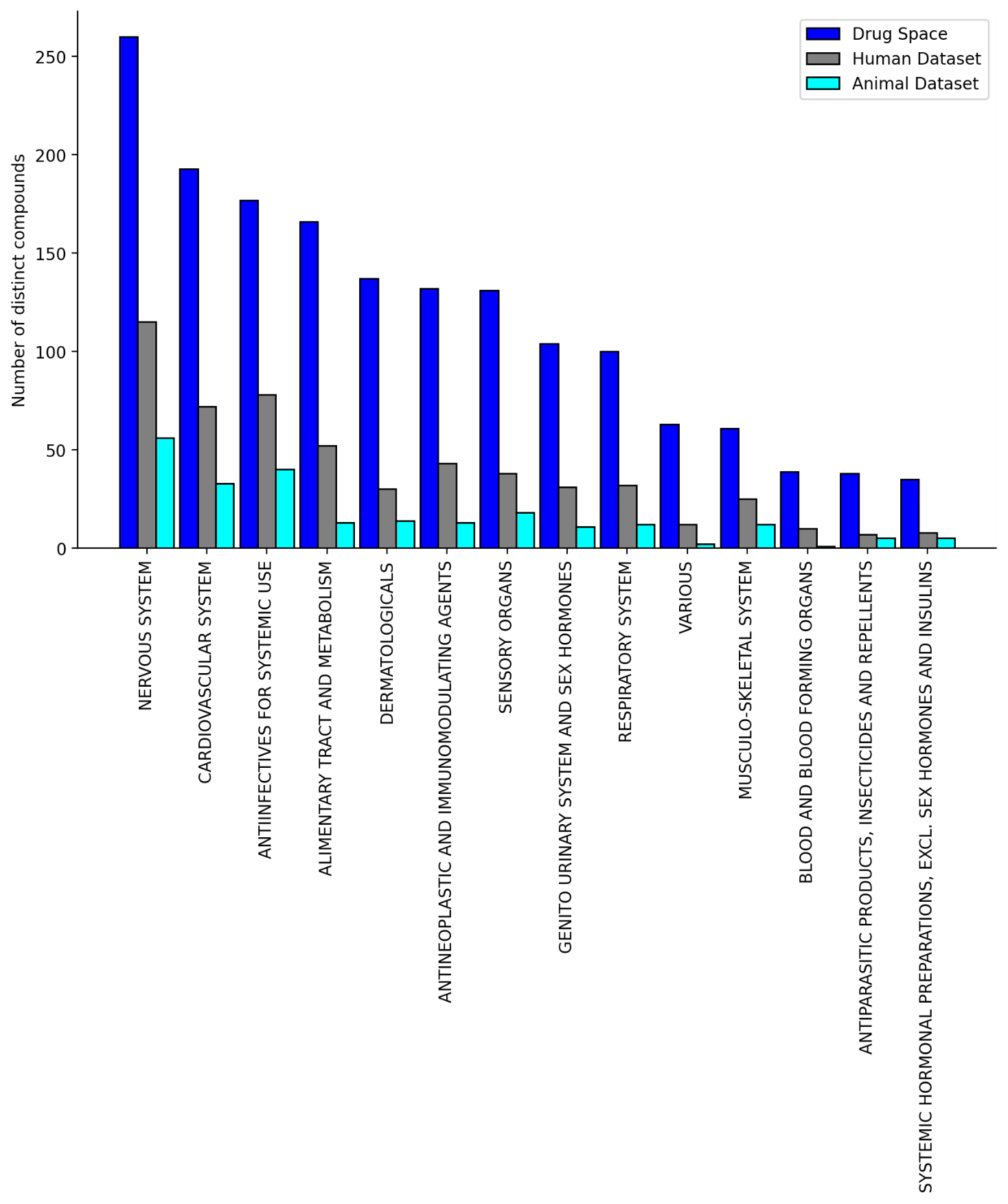

Figure S6: The human dataset and the animal dataset cover a broad range of ATC code distribution at the top level (for 553 out of 1283 compounds in the human dataset and for 235 out of 371 compounds in the animal dataset for which ATC annotations were available).

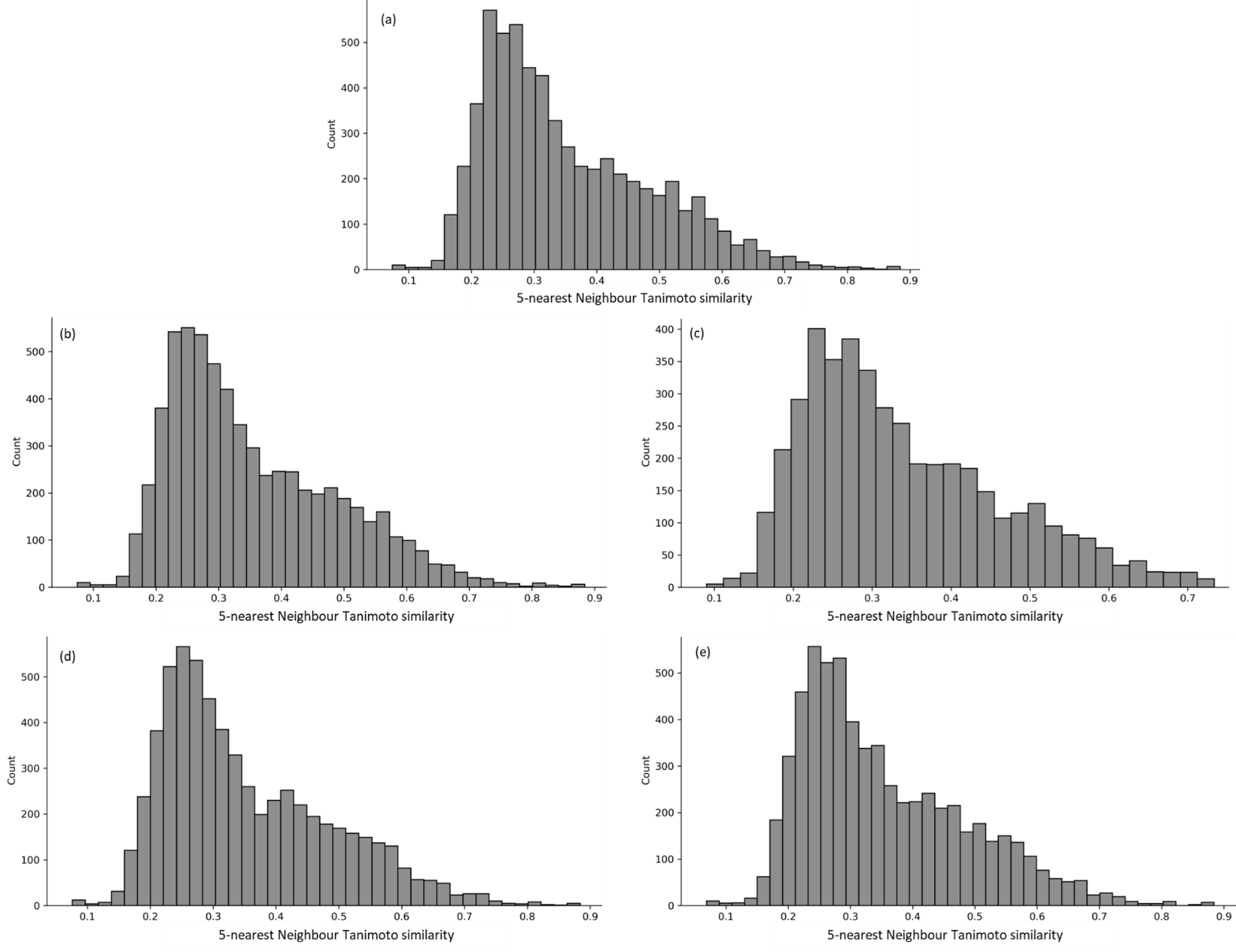

Figure S7: The mean 5 nearest neighbour similarity of all combinations of test compounds over the 25 test folds to respective training data in the nested cross validation for predicting the human PK parameters (a) Volume of distribution, (b) Clearance, (c) Fraction unbound in plasma, (d) Mean Residence Time, and (e) Half-life.

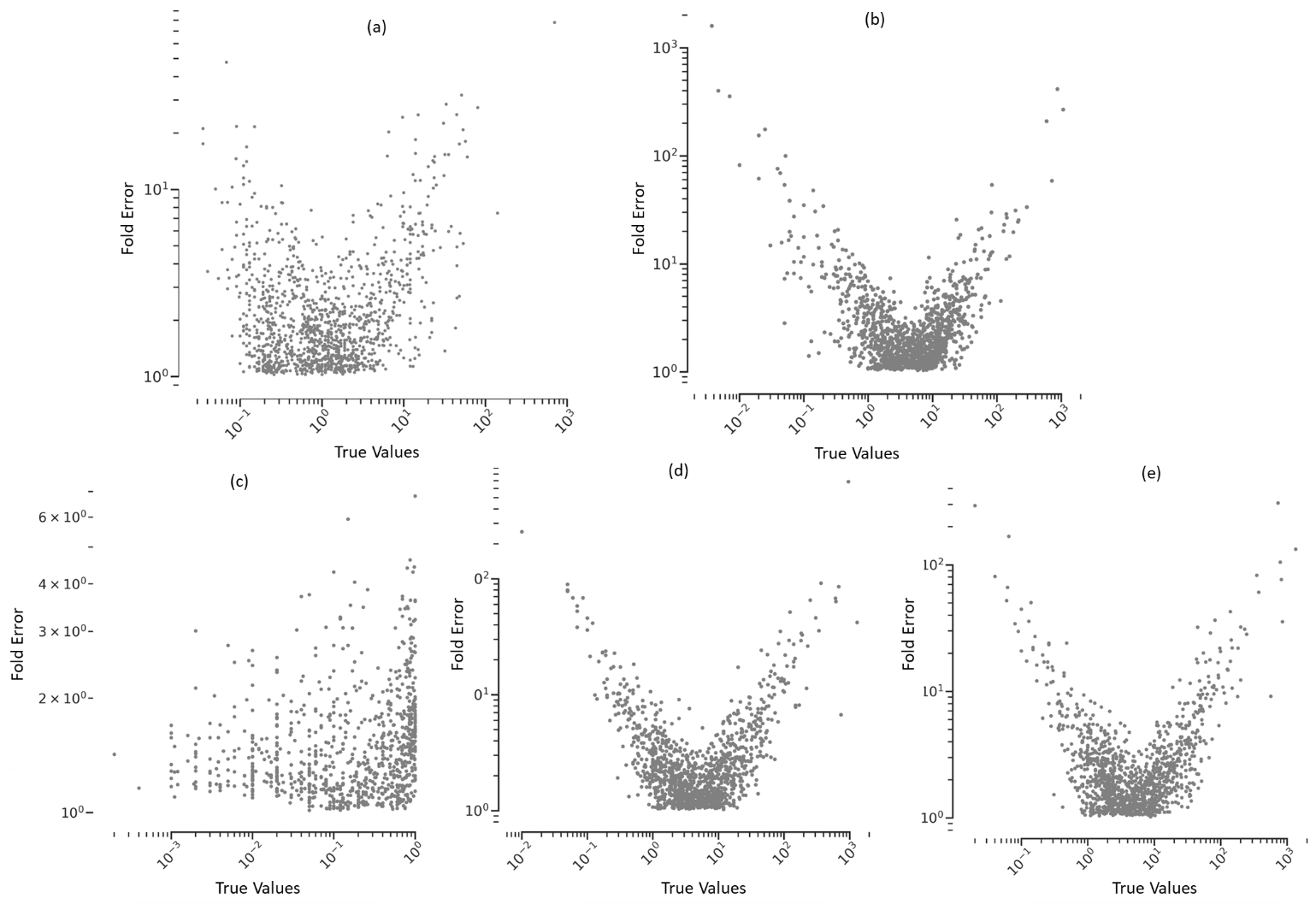

Figure S8: Comparison to fold error (of the predicted PK parameters) to the observed human PK parameters for all test compounds over the 25 test folds to respective training data in the nested cross validation for predicting the human PK parameters (a) VDss, (b) CL, (c) fu, (d) MRT and (e) t½. Fold errors tend to be large when the observed PK parameter value is further away from that of majority compounds.

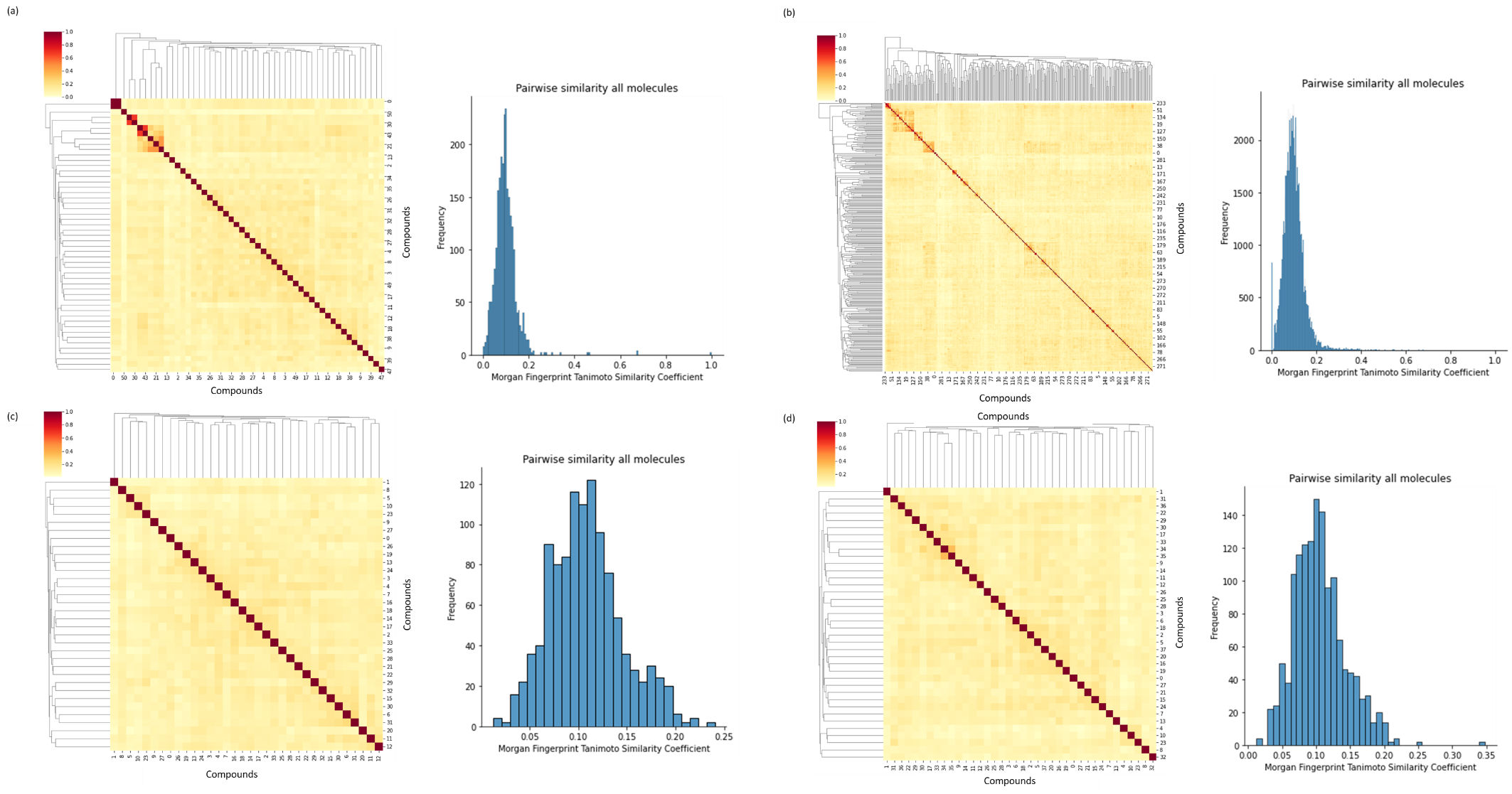

Figure S9: Pairwise Tanimoto similarity and the contour graph in the external test set using Morgan Fingerprints of radius 2 and 2048 nbits for (a) for 51 compounds for VDss, (b) 302 compounds for CL, (c) 34 compounds for fu and (d) 38 compounds for t½. Majority pairs of compounds (over 99%) are structurally diverse with Tanimoto similarity <0.30.
